# Hormones as adaptive control systems in juvenile fish

**DOI:** 10.1101/768689

**Authors:** Jacqueline Weidner, Camilla Håkonsrud Jensen, Jarl Giske, Sigrunn Eliassen, Christian Jørgensen

## Abstract

Growth is an important theme in many biological disciplines. Physiologists often relate growth rates to hormonal control of essential processes. Ecologists often study growth as function of gradients or combinations of environmental factors. Fewer studies have investigated the combined effects of environmental and hormonal control on growth. Here, we present an evolutionary optimization model of fish growth that combines internal regulation of growth by hormone levels with the external influence of food availability and predation risk. Hormones are represented by growth hormone, thyroid hormone and orexin functions. By studying a range from poor to rich environments, we find that the level of food availability in the environment results in different evolutionarily optimal strategies of hormone levels. With more food available, higher levels of hormones are optimal, resulting in higher food uptake and growth. By using this fitness-based approach we also find a consequence of evolutionary optimization of survival on optimal hormone use. Where foraging is risky, aerobic scope can be used strategically to increase the chance of escaping from predators. By comparing model results to empirical observations, many mechanisms can be recognized, for instance a change in pace-of-life due to resource availability, and reduced emphasis on reserves in more stable environments.

**Summary statement:** We combine physiological, environmental and evolutionary aspects of fish growth in a state-dependent model where the optimal regulation of growth and survival is achieved through hormonal regulation of behaviour.

## Introduction

It is a central aim of biology to understand how evolution has led to a specific organism design through natural selection. As Tinbergen (1963) pointed out, any trait can be understood both in terms of its mechanism and its evolution, and the philosopher Daniel Dennett (2017) has simplified this into two questions. If one for example is interested in fish growth, one may first ask “*How come* fish grow?” The discipline of physiology has excelled at answering this type of questions about underlying mechanisms, and has detailed triggers, pathways, intermediates, regulation, development, and function from the molecular level to that of the organism. There is another set of explanations for fish growth if one asks: *“What* do fish grow *for*?” “What for” questions are about the adaptive significance, about the effects a trait has on survival, growth, reproduction, and ultimately fitness. This evolutionary dimension introduces *purposiveness* to biology (Dennett 2017): a goal-directedness that goes beyond blind chains of causation like Hume’s billiard balls that crash into each other. Rather, processes occur to fill a purpose, to obtain some kind of aim, for example feedback processes that restore homeostasis, or drives or urges that ensure survival, growth, and reproduction. It must be emphasized that this is not an externally imposed or top-down purpose. It is a historic consequence of natural selection, where alleles with positive effects on survival and reproduction become more common in the gene pool, and their consequence is that organisms appear as goal-driven in their development, physiology, endocrinology, cognition, and behaviour (Andersen et al., 2016; Budaev et al., 2019; Giske et al., 2013).

“What for” questions have been addressed by evolutionary ecology, life history theory, and behavioural ecology, where empirical experiments and observations have often been inspired by theoretical considerations that have had one important limitation: they have typically ignored the proximate level of “how come” questions. This was epitomized by Alan Grafen as the phenotypic gambit, inspired by the chess move where one makes a sacrifice to gain a longer-term advantage (Grafen, 1984). The phenotypic gambit was a methodological tactic where one tossed away all the mechanistic detail and simply assumed unbounded phenotypic flexibility. Then and now, this was in many cases a necessary assumption to be able to answer “what for” questions. If models concluded that a trait had an adaptive advantage, the evolutionary ecologist would expect to see that trait to have evolved in real organisms in the wild. Any physiologist will immediately react to this as naïve and utterly unrealistic: Real traits originate from genes, are built through biochemistry, obey the laws of physics, and any information used must emerge from a sensory organ or use local molecules directly. The organisms that live today share many design features that have evolved precisely because they allow flexibility within the boundaries set by these constraints. Over time this has led to descendant lineages that were more likely to evolve to fill new niches and respond to new selection pressures. The combination of “how” and “what for” questions, thus, reveals insights that one of them alone could not give (Sinervo and Svensson, 1998). On the other hand, the traditional separation of mechanisms from the individual’s experienced selection pressures or ecological challenges, tears them out of a natural framework of constraints. It also builds on the assumption that selection pressures influence underlying mechanisms much less than the actual behaviour or adaptation they produce (Garland et al., 2016).

In this paper, we focus on one architectural design feature for control of the organism, its hormone system, and with a model we ask several questions that we believe are useful to stimulate thought both among physiologists and evolutionary ecologists. For example, are key hormone systems sufficient to enact the adaptive flexibility seen in growth across different environments? Are there ways in which we can conclude that the major hormone systems are adaptive? If we treat the model as a thought experiment with unlimited flexibility in hormone expression, will observed correlations emerge between environments and hormones? Between hormones? And with ontogeny? The model is about growth and related survival in juvenile fish, but more importantly it aims to show how one can overcome the phenotypic gambit, not only in the model specification, but hopefully also by helping scientists from the two disciplines in asking and answering questions together.

It can be instructive to compare our process-based model with other modelling approaches to better see the type of questions we can reach for. One type of well-known modelling tool in physiology are the dynamic energy budget models (DEB, (Kooijman, 2001; Kooijman, 1993; Nisbet et al., 2000; Zonneveld and Kooijman, 1989)). These follow resources and energy in great physiological detail from ingestion to growth and reproduction, and may provide good fit between predicted growth patterns and those observed in experiments and in the wild. One can describe DEB as “feed-forward bioenergetics”, where processes run as fast as resources or constraints allow. This perspective is similar to a combustion engine where the amount of gas fed into the carburettor determines the engine’s power and speed. Models of feed-forward bioenergetics are designed to question what happens to metabolic processes if more or less food is processed, when external conditions change, for example temperature, or when there are extra costs due to e.g. disease or reproduction. These are analogous to how fast a car would go if it is loaded heavy with passengers, if cooling is difficult on a particularly warm day, or if one of the spark plugs doesn’t fire.

In contrast our model optimizes survival through the juvenile phase, where the optimal growth rate emerges from the effects of growth on fitness. These may depend on the abundance of predators, food availability or duration of the growth season. Here, behaviour and physiology have to provide the resources required to achieve the target growth rate. This can be described as “by-demand bioenergetics”; a goal-driven control system that translates fitness incentives emerging in ecology into physiological responses that endow the phenotype with a performance to fulfil the set goal. This would be analogous to how hard the driver presses the gas pedal, which can depend on the speed limit, whether the driver is heading for the nearest hospital with a passenger about to give birth, or whether the passenger is a child who easily becomes car-sick. The car is a tool to achieve a goal in the driver’s mind, much like the physiology of an organism has potentials that can, if regulated appropriately, achieve fitness. So, while evolutionary ecology often seeks the optimal behavioural route to a goal, we here seek the optimal control mechanism along a given road.

There are several ways in which these control mechanisms can regulate and interfere with the individual’s bioenergetics. As the system is goal-driven a certain amount of energy has to be directed to mechanisms needed to achieve the goal. The process of allocation of limited resources towards competing uses (Fisher, 1930) is essential here. Also, as resources must be acquired before they can be distributed, the acquisition rate is of importance. Often models deal with either acquisition or allocation. Here we combine the two in one model organism and under one control system. In this way “by-demand bioenergetics” can drive the phenotype towards its goal by increasing the goal-directed energy supply through acquisition and allocation. Upregulating “by-demand bioenergetics” in such a way can push the organisms into a state of fast growth and early maturation. From an evolutionary point of view this would mean that life history changes from slow to fast.

Changes in growth rate are always accommodated by changes in other physiological, endocrinal and behavioural properties. This is due to the fact that mechanisms supporting growth have to be adapted to the new circumstances of fast growth, but also because of the cross-linking of mechanism and pleiotropic effects of hormones. Consequently, can a change in growth rate entail many other behavioural, physiological, endocrinal and life-history traits, which altogether form a suite of traits. This suite has been called pace-of-life syndrome (POLS, (Reale et al., 2010)). A special case of a fast life history is the “super” phenotype (Reznick et al., 2000) that makes use of rich environments by increasing its acquisition rate. “Super” phenotypes upregulate their energy-supply to all processes keeping allocation proportions constant. Thus, the whole phenotype is pushed into a highly energy-demanding but fast processing state.

To be specific about the goal-directness of growth in a proximate and mechanistic perspective, we treat the phenotype as having potential for a range of physiological rates, and focus on a simplified set of hormones as the control system. Because there are hundreds of hormones and associated signalling molecules in a typical fish or mammal, it was necessary to simplify to a level of complexity that is easier to grasp and analyse. We therefore first describe how we have interpreted the major regulatory routes that control growth in fish, and end up using three hormones and a neuropeptide that each play a specific role in our model. To a physiologist this simplification is most certainly incomplete as it definitely leaves out important elements, but our aim is to stimulate thinking, and we therefore ask the reader to follow us into this intermediate level of complexity. We now first describe how we have implemented our model, before we use the model to point to some interesting insights of the hormone system as adaptive, and ways forward to further bridging the proximate “how come” and the ultimate “what for” traditions in biology.

### Model

The model organism is a generalized juvenile fish, and we choose parameters mostly from Atlantic cod *(Gadus morhua)* which is a well-studied species. The model follows juvenile fish as they grow through a size window where they typically remain immature. During this juvenile phase we let internal mechanisms like metabolism and growth be regulated by two hormones, growth hormone and thyroid hormones, and the neuropeptide orexin. They determine growth, metabolic rate, and appetite, respectively, but importantly for the model they are also involved in trade-offs related to risk.

We use a state-dependent dynamic model (Clark and Mangel, 2000). This algorithm first optimizes a strategy that can be considered the evolutionary adaptation to a certain environment. In the case of this model the strategy is the optimal hormone levels for any combination of a fish’ size and energy reserves. When the optimal strategy has been found, we investigate this adaptation by simulating individuals that live in the given environment and use the calculated optimal policy, and we record its trajectory of growth, hormone expression, and individual states.

## Methods

### Simplifying the hormone systems for model implementation

The central challenge for our model organism is to grow (and survive) up to adult size. Although a high number of hormonal molecules and mechanisms are used to dynamically control physiology and behaviour in natural fish, we single out three clusters: growth, energy acquisition, and overall metabolism. When combined in a life history model, these also determine energy allocation to reserves. Below we describe the main hormones that work along these axes, and we call them “hormone functions” to distinguish them from real molecules. The main components of our mode are thus the growth hormone function, the orexin function, and the thyroid hormone function. Leptin also plays a role as it contains information about the individual’s energy reserves.

Decisions connected to growth influence the individual’s life history. For example, fast growth enables organisms to reach sexual maturity relatively early in their lives and start reproducing prior to conspecifics. Growth processes can make up a major part of energy use. The main endocrinal driver of growth in fish and mammals is growth hormone and its associated hormone cascade (Björnsson, 1997; Jönsson and Björnsson, 2002). Thus, in terms of “by-demand bioenergetics”, growth hormone drives the fish towards sizes at which they can mature and reproduce, implying that fitness considerations have set up an energy-demand that the organism needs to fulfil.

Part of the growth processes initiated by the secretion of growth hormone is the accretion of proteins and breakdown of lipids. Both processes influence the individual’s condition, and they increase metabolism. To maintain its condition, the individual must increase its energy uptake through foraging. Appetite and the initiation of feeding behaviour are very complex processes, comprising central nervous system and peripheral signals. An important group of neuropeptides are orexins, as they are produced in the hypothalamus, where signals on condition and energy-budget of the individual are integrated. Thus, orexins are the second step in the physiological response of the “by-demand bioenergetics” model, as they regulate the individual’s energy acquisition in order to fulfil the growth goal set by growth hormone.

To achieve growth, growth hormone as initiator and orexin as energy-suppliant are important factors influencing growth rate. Diving into growth mechanisms, there is another hormone and its associated cascade being ubiquitous for growth to happen: thyroid hormones. Hormones from the growth hormone cascade and the thyroid hormone cascade make up a complicated network in which they promote each other’s secretion, conversion, receptor activity and, in a chronological order, the developments of both cartilage and bone (Cabello and Wrutniak, 1989; Robson et al., 2002). Another reason for implementing a function on thyroid hormones is their regulating effect on metabolism (see below). On the one hand, an upregulated metabolism can be of advantage when energy is abundant. This would push the individual into a state of high energy turn-over. On the other hand, any increase in foraging exposes the individual to a trade-off between energy provisioning and foraging-related risk. The increased metabolism due to thyroid hormones can weaken this trade-off by allowing for faster metabolism and higher potential activity level, in turn causing higher escaping ability in case of a predator attack. In terms of the “by-demand bioenergetics” model the individual’s performance to fulfil the set growth goal is improved by higher energy turn-over and oxygen uptake rates when conditions allow.

Starting with empirical data on stimuli, hormone regulation, and effects, we now present the functions and mechanisms of these three clusters. Thereafter we will use this as background for the implementation in model code.

#### The Growth Hormone Function (GHF)

*Effects* Growth hormone (GH) is expressed throughout life. In humans, maximal secretion is seen during puberty, then decreasing with age (Vermeulen, 2002; Zadik et al., 1985). GH seems to affect metabolism and body composition (Velez et al., 2019; Vermeulen, 2002; Yang et al., 2018), but main effects are directed towards growth in bone (Nilsson et al., 2005; Robson et al., 2002) and muscles (Grossman et al., 1997). For fish, a relationship between GH levels and compensatory growth is suggested (Ali et al., 2003). To some extent GH also influences behaviour, either in a direct or indirect way (Jönsson and Björnsson, 2002). As growth rates can be constrained by environmental factors as food availability, GH levels and levels of its mediator IGF-1 should underlie seasonal fluctuations. Fluctuations, which might be stimulated by changes in photoperiod, have been observed in reindeer *(Rangifer tarandus)* (Suttie et al., 1991; Suttie et al., 1993) and Arctic char *(Salvelinus alpinus)* (Jørgensen and Johnsen, 2014).

*Axis* GH production is controlled by a hormonal cascade, the somatotrophic axis. On top, GH-releasing factor (GRF) and/or somatostatin (SRIF) are released by the hypothalamus upon environmental or peripheral stimuli. These regulate the anterior pituitary activity, which alters the rate of GH secretion. GH effects are mediated by IGF-1 in most tissues. Both GH and IGF-1 can affect mechanisms in target tissues (Gatford et al., 1998; Peter and Marchant, 1995).

*Stimuli* Through evolution the number of factors regulating GH release has decreased, while it is multifactorial in fish, regulation in mammals is mostly achieved by a “dual control system” (Gahete et al., 2009). The mammalian system consists of one main stimulator, growth hormone-releasing hormone (GHRH) and one main inhibitor, somatostatin (SRIF). Additional stimulators of minor importance are neuropeptide Y (NPY), ghrelin, exercise, and in some species leptin (Gahete et al., 2009; Hamrick and Ferrari, 2008; Kojima et al., 1999; Lanfranco et al., 2003). Leptin signals the current reserve size (Cammisotto and Bendayan, 2007), while ghrelin prepares the digestive tract for incoming food (Müller et al., 2015). In fish, a second main stimulator is pituitary adenylate cyclase activating polypeptide (PACAP). Additional weaker stimuli come from thyrotropin-releasing hormone (TRH), gonadotropin-releasing hormone (GnRH) and others. Leptin does not exert a direct stimulus in fish (Gahete et al., 2009).

Melatonin (Suttie et al., 1992; Suttie et al., 1991) regulates IGF-1 secretion. It is important to notice that one stimulus can have different effects on GH and IGF-1. This is for example the case in a study on fasted tilapia (*Oreochromis mossambicus),* where both body growth rates and body weight in males decreased due to fasting. IGF-1 levels correlated with growth rates, but GH levels were unchanged. A possible explanation is that available energy is used to cover basal metabolism first, while hormone levels are adapted to reduce or cease growth (Uchida et al., 2003). This is also the case for a diet experiment with Arctic char. Concentrations of growth hormone did not reflect changes in body weight, but IGF-1 concentrations did (Cameron et al., 2007). Unchanged or even elevated levels of GH can be part of a fasting response in which GH impels lipolysis and prevents protein degradation (Richmond et al., 2010).

Inhibition of GH is also exerted via IGF-1 in a long feedback loop, in both fish and mammals (Gahete etal.;, 2009).

#### The Orexin Function (OXF)

*Effects* Orexin is a neuropeptide known from humans (Kalamatianos et al., 2014; Oka et al., 2004; Tomasik et al., 2004), pigs (Kaminski et al., 2013), rats (Dube et al., 1999), and fish (Facciolo et al., 2010). There are two types of orexin, A and B, which have several effects, including feeding-related and behavioural effects (Cai et al., 2002; Rodgers et al., 2002). Orexin A stimulates foraging in goldfish *(Carassius auratus)* (Volkoff et al., 1999) and rats (Dube et al., 1999; Rodgers et al., 2000). Positive correlations between caloric demand and both orexin A and B exist for children (Tomasik et al., 2004). Observations of orexin A and B injected mice revealed no effect of orexin B on food intake, while orexin A increased food intake and metabolism (Lubkin and Stricker-Krongrad, 1998). One mechanism by which orexin can act on food intake is via regions in the brain as the arcuate nucleus (ARC) (Rodgers et al., 2002), where also leptin influences energetic processes in the body. It has also been suggested that foraging activity is increased by delaying satiety, as shown for low dose treatments in rats (Rodgers et al., 2000). Effects not related to feeding include a general arousal, reduced pain perception, increased locomotion etc. (Rodgers et al., 2002), and many of these can be seen as enabling for foraging. Despite of both orexins being present in a variety of organisms, the effect of orexin A on feeding behaviour seems to be much stronger than that of orexin B (Edwards et al., 1999; Haynes et al., 1999; Nakamachi et al., 2006; Sakurai et al., 1998).

*Stimuli* Factors influencing the secretion of orexin describe the body’s current state in terms of energy availability. A stimulating factor reported for rats is the fall in plasma glucose levels, eventually in combination with an empty stomach (Cai et al., 2002; Cai et al., 1999). However, a study on rats with insulin-induced fall in plasma glucose only showed an increase in hypothalamic orexin B (Cai et al., 2001). When energy is available to the organism, orexin secretion is inhibited. A signal of ingested food can be gastric distension (Cai et al., 1999). Leptin receptors have been found linked to orexin neurons in rodents and primates (Horvath et al., 1999) and may decrease the secretion of orexin in the hypothalamus (Kalra et al., 1999). Orexin A is believed to be part of a short-term response to ensure energy balance in the body (Cai et al., 1999; Rodgers et al., 2002).

Orexin effects in fish are similar to those in mammals (Matsuda et al., 2012) and they have been detected in several fish species (Miura et al., 2007; Nakamachi et al., 2006; Volkoff et al., 2003). Most experiments are done on goldfish (Penney and Volkoff, 2014), but also cavefish *(Astyanax fasciatus mexicanus)* show an increase of orexin A in relation to food intake (Penney and Volkoff, 2014). An interplay between orexin and ghrelin is suggested for foraging initialisation, in which ghrelin stimulates food intake and mediates orexin effects (Miura et al., 2007; Penney and Volkoff, 2014). Ghrelin is known from several fish species (Matsuda et al., 2011). In mammals, an increase in ghrelin-concentrations can be observed before food intake (Müller et al., 2015). In fish, it seems that patterns in ghrelin secretion are more species-specific. Several species show increases, as in mammals, but also decreasing concentrations are found (Jönsson, 2013; Penney and Volkoff, 2014; Rønnestad et al., 2017). Despite of differing mechanisms, it seems that the positive effect of ghrelin on foraging is similar across fish species.

#### The Thyroid Function (THF)

*Effects* In mammals and fish, thyroid hormones (TH) are major factors regulating metabolism and development. The hormones affect brain development (Di Liegro, 2008), metamorphosis (Youson et al., 1994) and, in combination with growth hormone, bone growth (Nilsson et al., 2005; Robson et al., 2002). Throughout life the basal metabolic rate is regulated by TH (Heilbronn et al., 2006; Herwig et al., 2008; Kitano et al., 2010; Webb, 2004). Due to their effect on metabolism they also play an important role in preparing organisms for seasons of low temperature and food availability (e.g. in red deer *(Cervus elaphus)* (Kuba et al., 2015), red knot *(Calidris canutus canutus)* (Jenni-Eiermann et al., 2002), reindeer (Bubenik et al., 1998) and white grouper *(Epinephelus aeneus)* (Abbas et al., 2012)). Consequently, some seasonal variation in circulating hormone levels can be detected. A reduction of up to 30% in basal metabolic rate in the absence of TH is documented for endotherms, and this reduction can be linked to thermogenesis (Heilbronn et al., 2006; Mullur et al., 2014; Silva, 2003). Non-thermogenic effects include the regulation of body weight and metabolism of triglycerides and carbohydrates (Mullur et al., 2014; Varghese and Oommen, 1999; Varghese et al., 2001). In both mammals and fish an impact on cardiac output is documented (Carr and Kranias, 2002; Little and Seebacher, 2014), and effects of TH on resting hearts has been shown in zebrafish *(Danio rerio)* (Little and Seebacher, 2014). As cardiac output contributes to maintain aerobic scope, TH also impacts the animal’s ability to sustain sufficient oxygen uptake under changing temperatures (Little and Seebacher, 2014).

*Axis* TH secretion depends on a hormone cascade sustaining relatively constant circulating hormone levels. On environmental or peripheral stimulation, thyroid releasing hormone (TRH) is secreted by neurons in the hypothalamus. In mammals, it promotes thyroid-stimulating hormone (TSH) release from the pituitary. In fish, the relation between TRH and TSH is not as clearly defined (Abbott and Volkoff, 2011; Chatterjee et al., 2001). In both mammals and fish, TSH acts on the thyroid gland, the actual place of TH production, which is stimulated to release TH into the blood. Those are mainly thyroxine (T_4_) but also triiodothyronine (T_3_), which differ in the number of their iodide ions (Han et al., 2004; Zoeller et al., 2007). Relatively constant hormone levels in the body are accomplished by negative feedbacks in the hormone cascade (Fekete and Lechan, 2014; Zoeller et al., 2007). TH are mainly eliminated from the blood by deiodination in the liver (Malik and Hodgson, 2002; Zoeller et al., 2007). The first deiodination-process forms the bioactive T_3_ from T_4_. There is also some evidence on the direct effect of TRH on feeding and locomotor activity (Abbott and Volkoff, 2011).

Target tissues, such as the brain, bones, and kidneys, contain different kinds of metabolic enzymes, deiodinases, to remove iodide from the hormones (Friesema et al., 1999; Miura et al., 2002). Biological inactive T_4_ has to be converted to T_3_ in order to have an effect on tissues (Zoeller et al., 2007). There are three deiodinases, which successively can remove iodide ions to form T_3_, T_2_, and T_1_ An inactive form called reverseT_3_ can also be produced (Zoeller et al., 2007). Although it seems that most studies concern the actions of T_3_, there is some evidence on effects of T_2_ (Lanni et al., 2001) and T_4_ (Robson et al., 2002).

*Stimuli* Several factors stimulating the release of TH have been identified, e.g. leptin (Abel et al., 2001; Herwig et al., 2008; Nilini et al., 2000) and insulin (Lartey et al., 2015). Leptin transfers information based on individual fat stores to the brain (Cammisotto and Bendayan, 2007), where the signal influences secretion of TRH positively (Fekete and Lechan, 2014). Inhibiting effects are known from stress (Silberman et al., 2002), exhaustive exercise (Hackney and Dobridge, 2009), and melatonin (Ikegami and Yoshimura, 2013; Ono et al., 2008).

#### Model Implementation

*GHF:* As our interest is in hormone strategies for growth, the growth hormone cascade is reduced to one variable in the model. This is a proxy for a fish’s IGF-1 blood plasma concentration and regulates the amount of energy drained from reserves and used for building all kinds of somatic structures, including bones. The complex hormonal network of ghrelin, leptin and the somatotrophic axis is resembled in the interaction of GH and current body states, notably energy reserves and satiety. In the model the axis, its effects, and stimuli are referred to as the growth hormone function (GHF) (Eales, 1988).

*OXF:* The orexin function (OXF) represents stimuli, hormone secretion, and effects of orexin as one value. For the model, only orexin A is regarded. To simplify its effects, the OXF only affects foraging behaviour in a positive manner. Foraging is assumed to include a series of other effects, such as arousal and increased locomotion, and in the model these are reflected in energetic foraging costs. Motivated from behavioural ecology, there comes a mortality cost with increasing foraging activity as looking for food involves potential encounters with predators. In the model we consider the longer term effect of the orexin function as a proxy for the mean orexin A concentration in the body during this period of time. Neither the effect of leptin nor ghrelin are modelled directly, but are integrated in the OXF hormonal mechanism.

*THF:* For the purpose of the model, a long-term effect of TH is of interest. Stress from predation, insulin and other factors that signal environmental or individual conditions on a short timescale are hence neglected. In the model the thyroid cascade is reduced to a simple factor resembling blood concentrations of bioactive T_3_. Negative feedbacks and elimination in order to receive relatively constant concentrations of TH in the body are disregarded; this is also done for the minor effect of T_2_ and T_4_. Effects of TH are reduced to an influence of thyroid on metabolism. Metabolism is regarded as the mean turnover of energy from food to reserves, soma, or activities. The influence of TH on metabolic mechanisms in the model is summarized in a positive linear correlation between TH concentration and standard metabolic rate (SMR). While this correlation is regarded as the cost of TH, a benefit comes with the positive linear correlation between TH and potential oxygen uptake, for example partly mediated through heart function. Increases in potential oxygen uptake through TH result in a greater free aerobic scope, which in turn contributes to higher escape rates in case of a predator attack. Non-metabolic processes as brain development or metamorphosis are not part of the model or the model fish’s life. As the “thyroid axis” in the model covers response to stimuli, the hormones themselves and their effects, it is called Thyroid hormone function (THF) (Eales, 1988).

### Model description

Hormones regulate physiological and behavioural processes, and these in turn achieve benefits and incur costs that may depend on the environmental conditions and the state of the organism. When we say we model hormones, it is therefore the *effects* of hormones that are in focus, in our case their consequences for growth and survival of juvenile fish. We first give the four central equations that describe growth and survival in our model, then detail the underlying processes. Throughout, capital letters are used for array variables that describe the organism and may change over time or with state, while lowercase is used for parameters that have a specific value (listed in Table 1). Greek letters denote the strategies, i.e. the hormone levels that the model optimizes.

The model characterizes fish body mass *W* [g] as being separated into two components, where the structural body mass *W*_structure_ [g] grows irreversibly. On top of that are the energy reserves *R* [J] that can be built or tapped, having an energy density *d*_reserves_ [J g^-1^]:

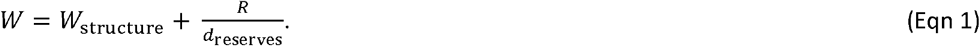

Growth Δ*W*_structure_[g week^-1^], the irreversible increase in structural body mass, depends on the level *γ* [ng ml^-1^] of the growth hormone function (GHF) relative to its maximum value *γ*_max_ [ng ml^-1^], current structural weight, and *k*_growth_ [week^-1^], which sets the upper limit for proportional increase in structural body mass per time step (weeks):

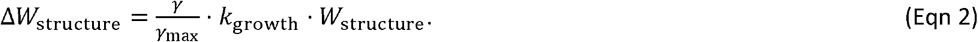

From the bioenergetics budget it follows that all energy taken up as food *I* [J min^-1^] is used for either metabolic processes *P* [J min^-1^] or to pay energetic costs of building tissues *C* [J min^-1^]. These new tissues include both new soma and changes in reserves.

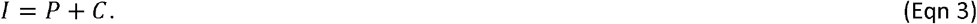

The details of *I, P*, and *C* are described in detail further down. Hormonally, *I* is controlled by the OXF, *C* by the GHF through tissue costs of growth, and *P* is influenced by the extra metabolic costs of expressing the THF.

The last central equation relates to surviva4 i I probability *S* [year^-1^], which is given by *S* = e^-M/52^ where *M* [year^-1^] is the total mortality rate compounded by several components:

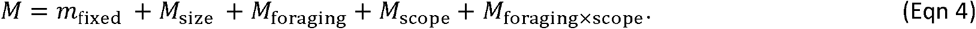

Here *m*_fixed_ is a constant irrespective of size, state, or strategy. *M*_size_ is a predation rate that declines with size. *M*_foraging_ is predation resulting from exposure while foraging. *M*_scope_ is increased vulnerability when the individual’s overall metabolic rate is close to its maximum aerobic capacity, because it is then harder to escape an attack. Similarly, *M*_foraging×scope_ is extra mortality when the individual exposes itself to predators while it is exhausted, which would put it in double jeopardy. The thyroid hormone function affects both *M*_scope_ and *M*_foraging×scope_.

Understanding the model requires that the equations above are interpreted in light of three key trade-offs that we describe here and give details and equations for further down.

First, the energy requirement of growth and everything else has to be met by foraging for food, which involves taking some level of extra risk (Krause and Godin, 1996; Lima and Dill, 1990; Sih, 1992). A resting fish often seeks safety in a shelter but needs to leave this to seek habitats where prey, and most often predators, are more common. Acquisition of more food thus involves more encounters with predators, and when food is scarce the fish needs to search for longer and expose itself more to forage the same amount.

Second, aquatic breathing is rapidly limited by surface-to-volume ratios and gas diffusion, even for small organisms. Although respiratory organs such as gills have evolved to overcome these constraints, there are physical limits to permissible total metabolic rate (Priede, 1985). Maximum aerobic capacity is often measured on fish that swim in respirometers, but digestion and growth are also variable processes that contribute to total metabolic rate. When the overall level of metabolic processes requires a lot of oxygen, the fish is quickly exhausted and therefore less efficient at evading predators should it encounter one.

Third, a trade-off that has received less attention is how spending energy can help an organism to manage, mitigate, or reduce risk. It is known that immune systems incur energetic costs, and that the optimal level of immune function depends on energetic status, the risk of infections, and availability of resources. Here we use thyroid regulation of metabolic level to achieve a similar exchange between energy and risk. The model assumes metabolic level can be upregulated by thyroid at an energetic cost (subject to trade-off 1), and the extra metabolic capacity is modelled as an elevated aerobic scope (alleviating trade-off 2). Consequently, the model allows metabolic rate to vary systematically between ecological settings.

We use a state-dependent model to find the optimal hormonal control of acquisition and allocation of energy. This type of mechanistic model finds the evolutionary endpoint (beyond which further changes cannot improve fitness) for a given environment. The model first uses dynamic programming (Clark and Mangel, 2000; Houston and McNamara, 1999) to find the optimal hormone expression, the strategy, for each combination of the individual’s states. The individual states included are the body length of the fish and its energy reserves. Thereafter, an individual that makes use of the optimal strategy according to its current individual state is simulated. We record its trajectory of growth, physiology, behaviour, and risk-taking to quantify and analyse effects. The model optimizes the state-dependent trajectory of the three hormones (GHF, OXF, and THF) by maximizing juvenile survival between 10 cm and 30 cm body length. The time steps are set to one week to represent typical dynamics of hormone levels and growth processes, which means that more rapid processes like behaviours are not modelled in minute-to-minute detail but for their cumulative effects at a weekly scale. The model describes growth of a juvenile fish in environments with constant food availability, and we compare several different environments in our analyses.

### Energy budgets and metabolic rate

The total metabolic rate *P* [J min^-1^] is the sum of all respiratory processes, all with unit joules:

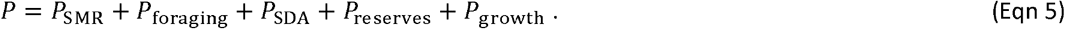

Here *P*_SMR_ [J min^-1^] is the standard metabolic rate, *P*_fomging_ [J min ^-1^] the swimming cost of foraging behaviour, *P*_SDA_ [J min^-1^] the cost of digestion and energy uptake (SDA) until the resources are available in the bloodstream, and *P*_reserves_ [J min^-1^] and *P*_growth_ [J min^-1^] the metabolic costs of converting between resources in the bloodstream and reserve and structural tissue, respectively.

On top of that, the organism uses its digested resources for incorporation as new structural tissue (*C*_growth_ [J]) or by adding to or using from energy reserves (Δ*R* [J]). The net rate *C* [J min^-1^] of such incorporation of energy into tissue is thus:

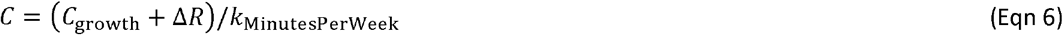

Note that while *P* and *C* both contribute to the individual’s energy budget (Eqn 3), only *P* uses oxygen through aerobic respiration (Eqn 24).

The basis, standard metabolic rate (SMR), scales allometrically with body mass as the fish grow from juvenile to adult size. Other contributors to an individual’s overall metabolic rate are factors like locomotion, digestion, and growth, and many of these may change with ontogeny (Mozsar et al., 2015).

The model uses variants of SMR in several ways. For what is measured experimentally as SMR and that we refer to as *P*_SMR_ is the standard oxygen consumption of the organism’s total body mass as it is affected by the level of the thyroid hormone function. The standard level of SMR at a mean level of THF expression is:

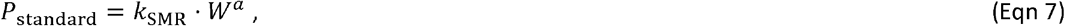

Where *k*_SMR_ has unit [J min^-1^ g^-a^], *P*_standard_ can be up-xor downregulated under the influence of THF, modelled as the concentration *τ* [ng I^1^] and relatively to a maximum concentration *τ*_max_ [ng ml^-1^]:

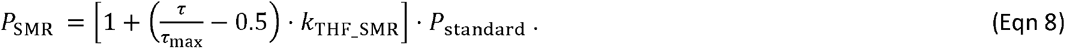

Here *k*_THF_SMR_ determines the strength of the effect of THF on metabolic rate, or in other words, the energetic cost of upregulating the scope for metabolic activity. It is *P*_SMR_ that enters the individual’s metabolic rate (Eqn 5).

When we model food intake as a multiple of SMR, it is unlikely that a chubby individual has higher foraging success per time and energy investment compared to a leaner fish, so we scale food intake with *P*_structure_, a measure of SMR calculated from the lean body mass only and not affected by THF:

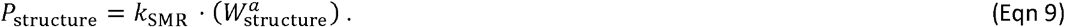

### Foraging and digestion

Energy from foraging is ultimately used to drive all energy-dependent processes in the organism. We model foraging as controlled by appetite through the orexin function where the relative concentration of 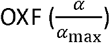 is proportional to the target intake rate *I* of the individual.

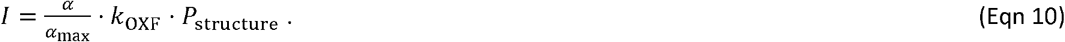

Intake *I* [J min^-1^] is defined as metabolizable energy absorbed by the gut; urinary and fecal loss of energy are implicitly included in the dimensionless coefficient *k*_OXF_ (Bureau et al., 2003). Here *P*_structure_ is a standardized metabolic rate of the lean body mass, explained in Eqn 9 above, used because it is unrealistic that having large reserves contributes to more efficient foraging.

The foraging behaviour *B*_foraging_ [dimensionless, given in multiples of *P*_structure_] required to meet the energetic demand depends on food availability in the environment. We first rescale foraging intake to multiples of SMR and then assume that food is quicker and safer to find in rich food environments *E* [dimensionless]:

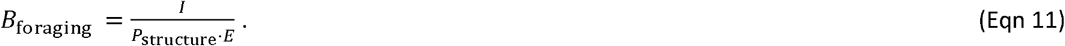

The cost of foraging activity (*P*_behaviour_) is proportional to foraging activity and SMR with a coefficient *k*_foraging_ [dimensionless]. Physical activity during foraging requires moving the whole body, including soma and reserves, so SMR is based on total weight.

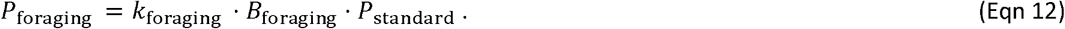

Food eaten is processed by the digestive system and taken up into the bloodstream. Specific dynamic action SDA (*P*_SDA_), representing the cost of digestion, is the product of intake and a constant *k*_SDA_ [dimensionless].

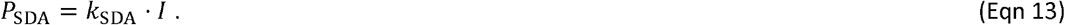

### Growth and reserves

Structural weight (*W*_structure_) is calculated based on length *L* [cm] using Fulton’s condition factor for lean fish (*k*_Fultons_min_, [g cm^-1^])

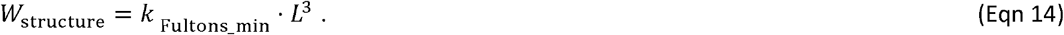

Likewise, maximum storage depends on body size and is calculated from the difference between maximum (*k*_Fultons_max_, [g cm^-1^]) and lean condition factor, and the energy density of the reserves (*d*_reserves’_ [J g^-^]):

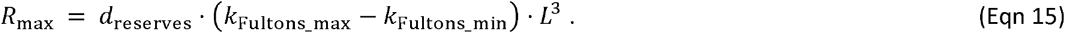

The cost of structural growth *C*_growth_ follows directly from the amount of new tissue produced (Eqn 2) and the somatic energy density *d*_structure_ [J g^-1^]:

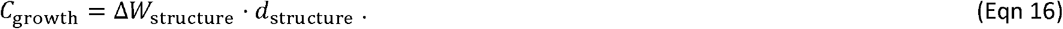

While reserves may vary in size, the model assumes that structural growth is irreversible (*C*_growth_ ≥ 0). A breakdown of soma, e.g. muscle tissue during starvation as seen in nature, is thus restricted to the part included in the reserves.

To meet the requirements of different tissues, nutrients have to be converted, and conversion of metabolites comes with a cost. When storing energy, processing of nutrients into storage molecules is based on a conversion efficiency *k*_conversion_reserves_ [dimensionless]. The model assumes this conversion to be biochemical processes that requires oxygen and therefore will contribute to overall metabolic rate:

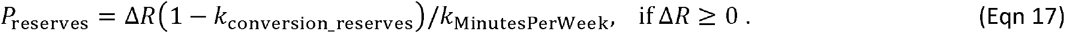

If energetic expenses exceed the energy available from digestion, reserves have to be drained. Then a conversion cost has to be paid for making those reserves accessible:

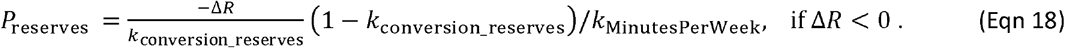

In the case of growth, metabolites are drawn from reserves and converted into building blocks. The cost *P*_growth_ of conversion into growth is also calculated using a conversion efficiency parameter *k1*_conversion_growth_ [dimensionless].

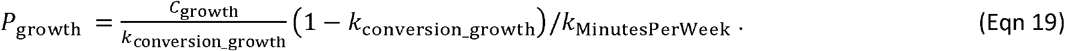

### Aerobic scope

The maximum rate of oxygen uptake has to accommodate all oxygen-dependent processes such as digestion, locomotion, foraging, conversion of energy, and other metabolic activities (Fry, 1971). We refer to the unused surplus as the free aerobic scope (Holt and Jørgensen, 2015).

We calculate potential oxygen uptake *A*_standard_ [J min^-1^] following Claireaux et al. (2000) as an allometric function with exponent *b* < 1. Because it is unrealistic that variations in reserve size affect an individual’s capacity for oxygen uptake, we base calculations of aerobic scope on the structural body mass only:

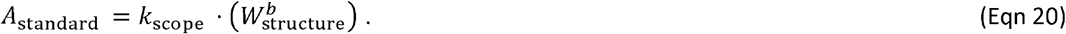

Here *k*_scope_ has unit J min^-1^ g^-*b*^.

A key assumption of our model is that the thyroid hormone function THF increases aerobic scope through increasing capacity for oxygen uptake, thus permitting higher levels of metabolic processes, but at a cost on SMR (Eqn 8):

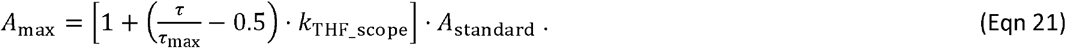

Here *k*_THF_scope_ [dimensionless] sets the strength of the effect of THF on increased scope.

### Food availability

Across model runs we vary food availability, implemented as the factor *E* [dimensionless]. When food availability is good (high *E*), less foraging activity is required to obtain the given amount of resources (Eqn 11). Contrary, when *E* is low, the individual needs more time to gather the amount of food it aims for. Consequently, *E*, through B_foraging, determines the exposure to predators in Eqn 23, and the energetic cost of foraging in Eqn 12. In this version of the model, there is no stochasticity influencing foraging success.

### Mortality rates

Mortality is decompounded into discrete risk factors (Eqn 4) that through separate trade-offs contribute to an individual’s risk of being depredated or otherwise die (extended from Holt and Jørgensen (2014)). The first is a constant component *m*_fixed_ that represents death due to causes that are independent of the individual’s state or behaviour, e.g. some types of disease. Second is size-dependent mortality, with reduced risk of mortality with larger body size, as is both observed (Gislason et al., 2010; Peterson and Wroblewski, 1984) and resulting from the size-structure of marine food webs and scaling relationships (Brown et al., 2004). We model this as an allometric relationship with a negative exponent:

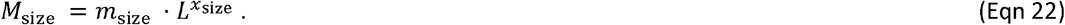

The next mortality component reflects the well-known trade-off between risk of predation and foraging intensity (e.g., Lima, 1998).The model assumes that individuals expose themselves to predation risk while foraging, and that this risk accelerates with increasing foraging because the safest habitats and time periods are assumed exploited first:

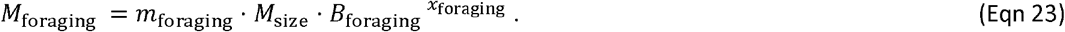

For this and the risk components below, it is assumed that predation is the ultimate cause for death and therefore that the risk declines with size in the same way as the size-dependent predation mortality.

The final two components relate to oxygen use and aerobic scope, i.e. the difference between maximum oxygen uptake and actual rate of oxygen use. Fleeing from predators demands burst swimming, which is achieved anaerobically by white muscle (Johnston, 1981; Rome et al., 1988; Weber et al., 2016). Recovery is aerobic and faster if there is free aerobic scope to provide abundant oxygen (Killen et al., 2014; Marras et al., 2010), thus preparing the individual for a repeated attack or the next encounter. We model this based on the ratio between used and available oxygen, raised to a power to describe how predation risk increases rapidly as maximum oxygen uptake is approached or even temporarily exceeded:

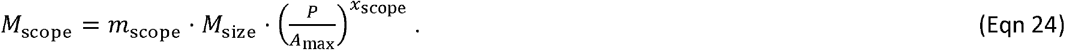

The model finally assumes that it is particularly risky for an individual to expose itself (high *M*_foraging_) when oxygen use is high (high *M*_scope_) because attacks would be frequent and recovery at the same time slow:

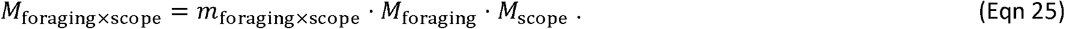

The mortality rates stemming from each risk factor are then summed (Eqn 4) and survival per time step given as *S* = e^-*M*^.

### Implementation

The model follows juvenile fish as they grow from 10 cm to 30 cm body length. Optimal solution is found for each combination of individual states length (21 steps) and reserves (10 steps). Discretization 160 steps for each hormone. Time step 1 week. Sufficient time horizon, normally 200 weeks.

### Parameterization

Parameters used in the model were chosen from different fish species to create a generalized, juvenile fish. Many of the studies used were performed on cod, which makes cod the fish most similar to the model fish.

For orexin A no studies on hormone concentrations in fish are known. In this case measurements on mammals were used.

The water temperature is constant at 5°C and water is saturated with oxygen.

Energy density for reserves is chosen to be 5 000 J/g. This is based on a calculation of mean protein and fat contents in storage tissues. A fish of 750 g serves as template. Energy density is based on the weight of liver and white muscle tissue and their proportional content of fat and proteins. For proteins, the weight of cellular water is taken into account.

Since growth requires development of more specialized tissue than storing molecules in reserves, the conversion efficiency for growth is lower than for reserves.

Fulton’s condition factors for fish with full reserves (*k*_Fultons_max_) and depleted reserves were chosen following a study on cod (Lambert and Dutil, 1997b).

Variables used in calculations of SMR (*k*_SMR_, *a*) are based on Clarke and Johnston (1999), Mozsar et al. (2015) and Pangle and Sutton (2005) accounting for the resting metabolic rate of a general teleost fish. In line with earlier models built on a similar bioenergetics template (e.g. Jørgensen and Fiksen 2010), we use a scaling exponent a=0.7 which is within the range of intraspecific scaling exponents for in teleosts (Killen et al., 2007). Also, studies show that there is a great variation for scaling exponents in animals and the value chosen here is in the range of this variation (Holdway and Beamish, 1984; Kjesbu et al., 1991; Lambert and Dutil, 1997a). Units are converted to fit the model.

The coefficient *k*_scope_ used in calculations is derived from a study on cod (Claireaux et al., 2000). The scaling exponent for aerobic scope (*b*) is chosen in accordance to SMR scaling (Holt and Jørgensen, 2014).

### Hormone Concentrations

Concentrations of IGF-1 are given in ng/ml blood plasma and range from 0 to 200. In experiments with tilapia concentrations of 70 – 120 ng/ml plasma were measured (Uchida et al., 2003). A study on Arctic char revealed concentration up to approximately 250 ng/ml plasma (Cameron et al., 2007).

Orexin A has been detected in ranges up to roughly 350 pg/ml porcine blood plasma (Kaminski et al., 2013). A range assumed to be normal for adult men and women (Oka et al., 2004). The range is higher for children, where measurements up to roughly 1300 pg/ml have been observed (Tomasik et al., 2004). For the model orexin A adopts a range up to 2000 pg/ml blood plasma. Its existence and function in fish has mainly been documented in goldfish (Abbott and Volkoff, 2011; Hoskins et al., 2008; Volkoff et al., 1999) and zebrafish (Matsuda et al., 2012).

Concentrations of T_3_ are given in ng/ml of blood plasma and range from 0 to 5. The range is chosen according to measurements on teleosts like one-year old rainbow trout *(Oncorhynchus mykiss)* (Eales, 1988), *Anabas testudineus* (Varghese and Oommen, 1999; Varghese et al., 2001) and chum salmon (*Oncorhynchus keta)* (Tagawa et al., 1994) revealing concentrations up to roughly 4.5 ng/ml plasma for normal individuals.

## Results

During the fish’s growth phase, the optimal strategy for the hormone profile changes, resulting in a near-linear length growth and decreased mortality rate over time (Fig. 2). While energy gain and oxygen budgets are relatively stable per unit body mass, mortality decreases with size. The optimal level of GHF falls throughout the growth phase (Fig. 2A), but as their effect is relative to body size, the resulting growth in length is near-linear (Fig. 2D).

**Figure 1.**
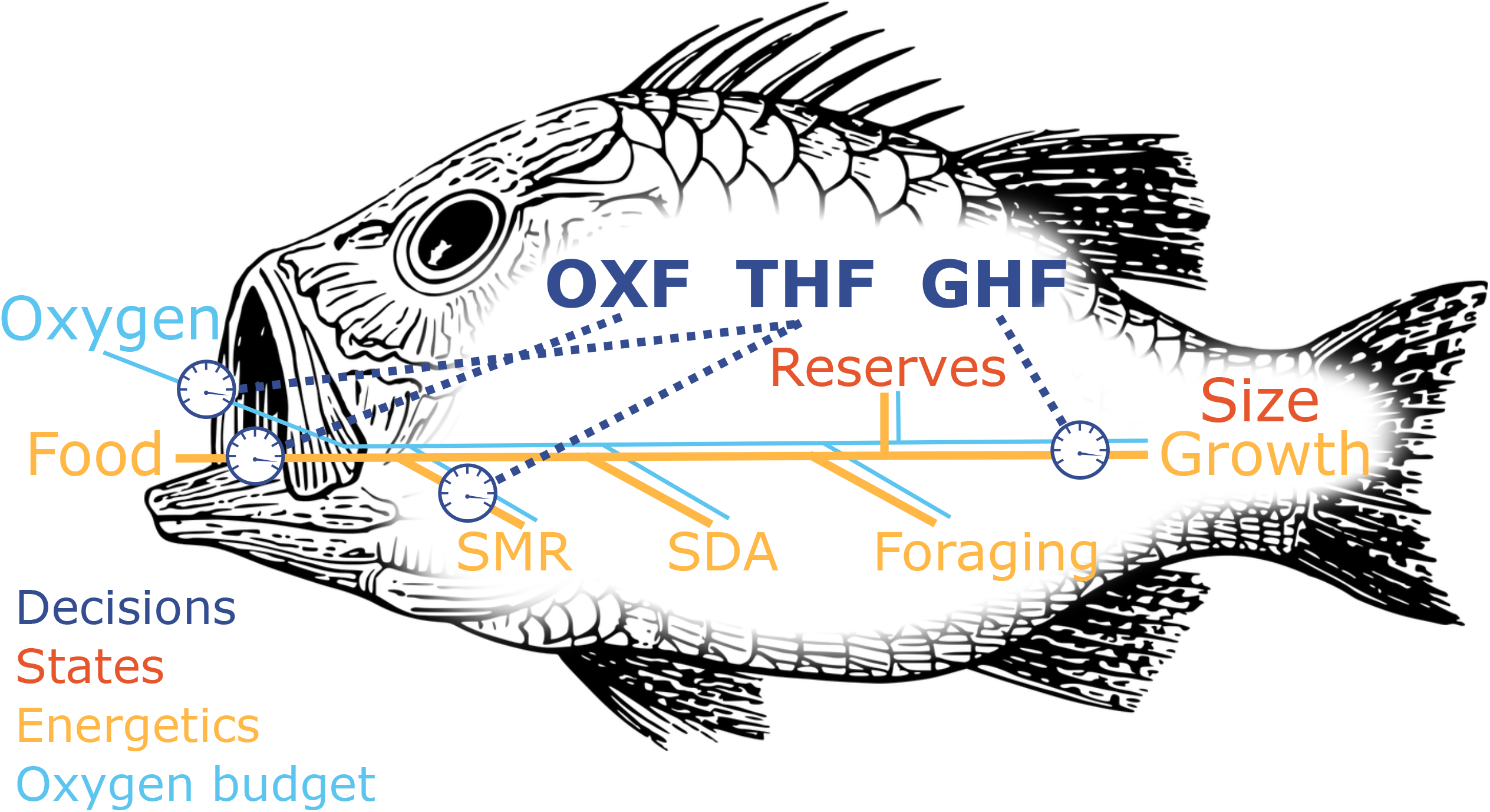
Energetics and endocrinology of the model organism. Energy from food is made accessible for the body by digestion (SDA). This energy is then used in metabolism to maintain life-supporting metabolic pathways (SMR) and supply the organism with oxygen. Also activities like foraging use energy. The surplus is stored in reserves. Hormonal regulation determines the foraging intensity (OXF), in- or decreases of metabolism rates (oxygen uptake & SMR), and the allocation of energy to growth (GHF). Throughout the simulation decisions regarding hormone levels are based on the two states of the fish – reserve and body size.

**Figure 2.**
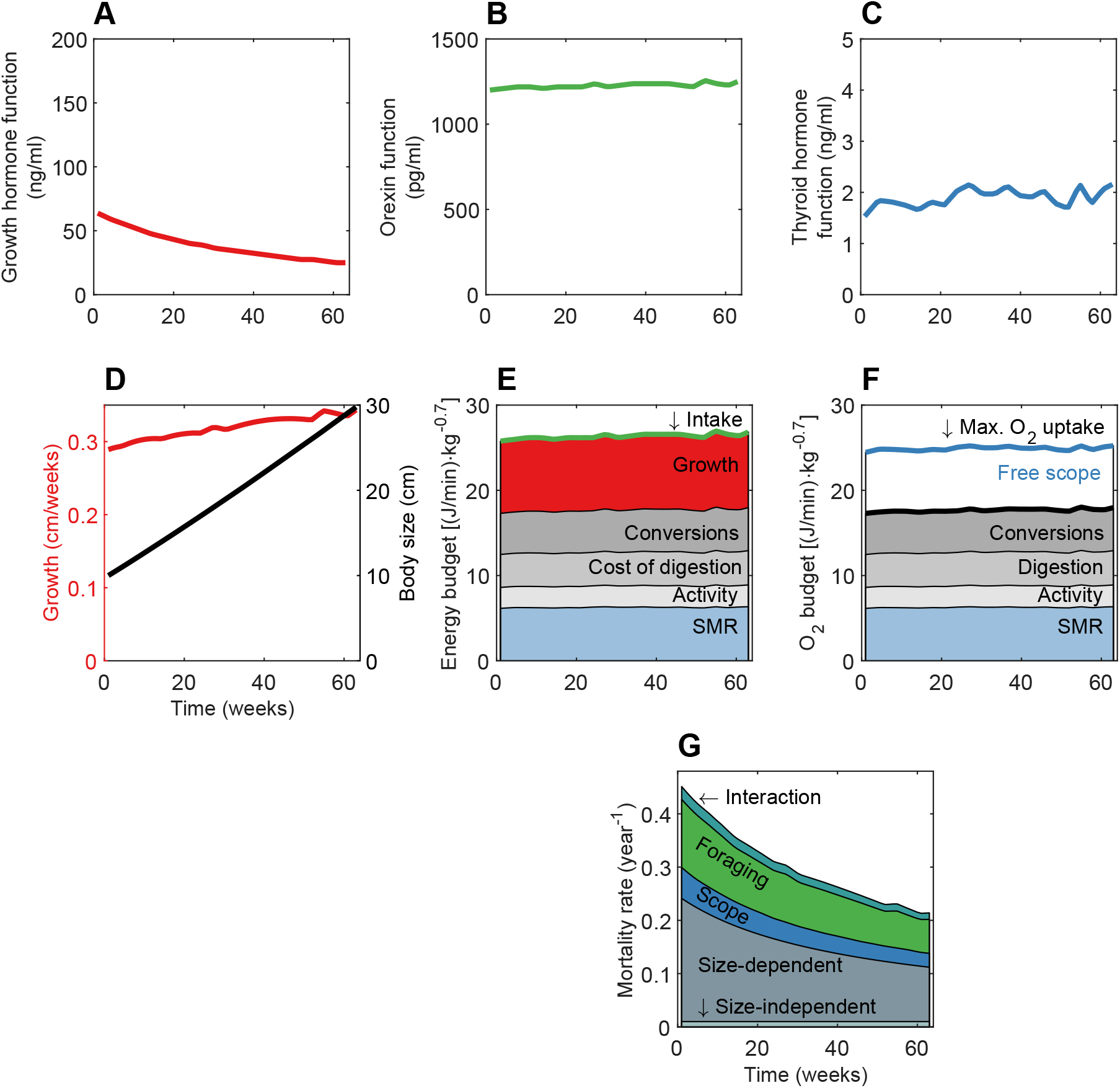
Endocrine regulation, energy and oxygen budget, mortality and growth of juvenile fish in a stable food environment. The simulation starts when the fish is 10 cm and ends at 30 cm, with the x-axis giving time (in weeks since 10 cm) in all panels. In A) the growth hormone function, B) orexin function, and C) thyroid hormone function, is given as a function of time. D) Weekly growth and accumulated body mass, E-G) energy budget, oxygen budget and mortality rate, respectively.

The optimal level of OXF (green) is relatively constant through the growth phase (Fig. 2B), which gives a stable food intake rate per body mass. Energy from feeding is allocated to SMR, SDA, soma, metabolic processes involved in conversion of food to reserves and growth, and the activity associate with searching for food (Fig. 2E). Since the food environment is not changing over time, the fish does not benefit from storing energy in reserves, but rather allocates all somatic investments in structural growth (Fig. 2E).

There is some variation seen in the levels of THF over the growth period for the fish (Fig. 2C). This variation is too small to have a visible effect on SMR or maximum oxygen uptake per metabolic mass (Fig. 2E & F). However, both SMR and maximum oxygen uptake for the individual increase due to increases in total body mass (not shown).

The instantaneous mortality rate decreases during development (Fig. 2G), mainly because size-dependent mortality (grey area, Fig. 2G) is smaller for larger fish (Eqn 22). Foraging mortality (Eqn 23), scope-related (Eqn 24), and active-while-vulnerable mortality components (Eqn 25) also drop. Foraging activity and free scope are relatively constant, hence changes in these mortality components are mainly due to lower predation risk with increasing size.

If we study how the optimal hormone strategies change along an environmental gradient that varies in food availability, we see that the levels of OXF, GHF, and in particular THF are higher in environments with more abundant food (Fig. 3A). Individuals in rich food environments grow faster, and have higher oxygen-uptake and better survival probabilities. Faster juvenile growth requires increased energy intake, which results in higher SDA and conversion-related costs. Oxygen requirements also increase, which selects for higher THF levels that increases maximum oxygen uptake and secures free scope (Fig. 3C). THF also upregulates SMR, hence the optimal hormone level depends on the availability of energy in the environments and costs in terms of energy and mortality that come with gathering food. The energy allocation trade-off, between investments in maintenance and survival on the one hand, and growth on the other, changes with food availability. Throughout the growth phase this trade-off is influenced by THF, deducting energy to support a higher metabolic rate that in turn increases escapement probability from predators. As energy is more accessible when food abundance is higher, activity costs are unchanged even when intake increases (Fig. 3B). Due to higher hormone levels, fish in habitats with high food availability have higher growth rates, intake, and SMR (Fig. 3).

**Figure 3.**
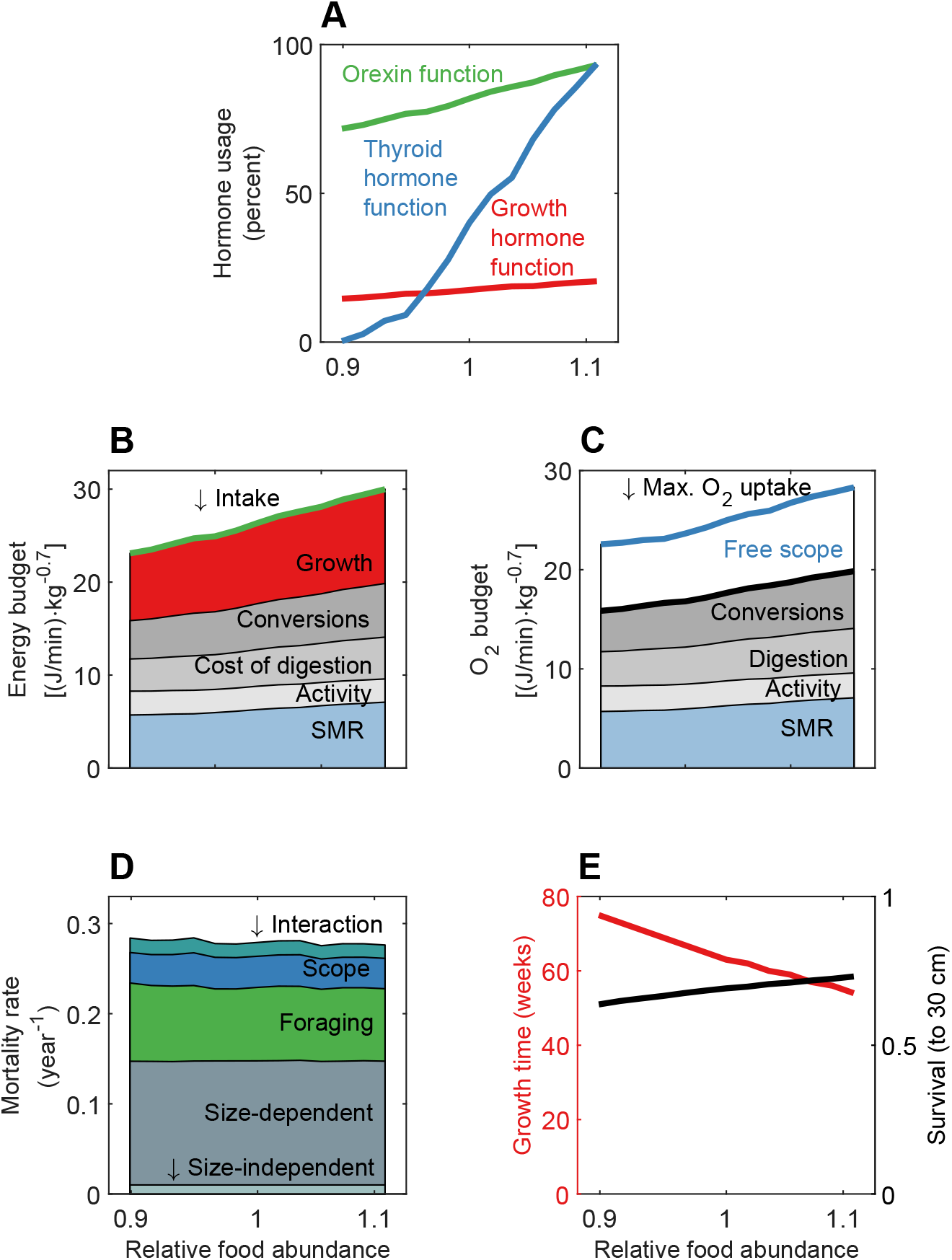
Environmental impact on hormone levels, energy and oxygen budgets, survival and generation time. The x-axis is the same in all panels, with a gradual increase in food abundance relative to the average food environment used in Fig. 2. Simulations of fish in 13 food environments are compared, at individual length around 20 cm. A) Hormone levels B) Energetic costs from growth and metabolism. C) Free scope, as the difference between maximum oxygen uptake and the sum of processes consuming oxygen. D) Five different components contribute to mortality. E) Growth time and survival over the entire juvenile life phase of the fish.

Comparing oxygen budgets (Fig. 3B), we see a slight increase in free scope from the poorest to the richest food environment. THF enables the organism to increase its free scope despite higher oxygen use, thus permitting higher growth and foraging through the other hormones. Oxygen used for preparing metabolites for new soma reduces free scope, while THF works against this process by elevating maximum oxygen uptake.

Simplified, GHF sets energetic needs, OXF meets the needs by determining foraging activity and providing metabolites for growth. The increased energy turnover has to be supported by THF, regulating maximum oxygen uptake to reduce mortality rate when energy is readily accessible and high turnover desirable (Fig. 3D).

Adaptations in hormone levels cause fish in rich environments to have a shorter juvenile phase (Fig. 3E). Despite similar instantaneous mortality rates (Fig. 3D), the probability of surviving to the end of the growth phase differs substantially between food environments because the duration of the growth phase is longer when food is scarcer.

## Discussion

Most evolutionary optimization models of animal growth and survival focus on behaviour, size, or other phenotypic traits while the internal regulatory processes are often ignored (Fawcett et al., 2014; Grafen, 1984). For fish, this includes social behaviour (Rountree and Sedberry, 2009; van der Post and Semmann, 2011), diel vertical migration (Burrows, 1994), and habitat choice (Fiksen et al., 1995; Kirby et al., 2000), but see Salzman et al. (2018). Here we take the opposite perspective, and study optimal internal regulation by hormone systems for animals that cannot choose their external environment. Obviously, most animals can do both at the same time, and habitat selection can have direct impact on the physiological needs and priorities of the animal (Elton, 1927). But by removing the movement options in this model, we can isolate how internal mechanisms can be used to optimize trajectories of growth and mortality risk. We found variation in optimal hormone levels across different food environments and throughout ontogeny. We modelled adaptive evolution in three hormone functions, where the growth hormone function (GHF) sets the fitness-optimizing growth rate, the orexin function (OXF) provides the required resources through appetite control and foraging, while the thyroid hormone function (THF) adjusts trade-offs between bioenergetics and survival. The effects of the hormonal control are evident in growth patterns, energy allocation, oxygen budget, activity levels, and in survival.

Increased food availability enables organisms to grow faster, which is achieved by speeding up metabolism to accommodate increased physical and biochemical activity. Model fish adapted to high food availability by having higher optimal concentrations of GHF and THF than those adapted to food-restricted habitats (Fig. 3). Empirical studies testing for changes in hormone concentrations in relation to diet quantity focus on short-time experiments, often with feeding – starvation – refeeding cycles. Similar to the predictions of the model, these generally find a positive correlation between hormone concentrations in plasma and the amount of food eaten by the fish (Lescroart et al., 1998; MacKenzie et al., 1998; Power et al., 2000; Toguyeni et al., 1996; Van der Geyten et al., 1998) or mammal (Herwig et al., 2008; Lartey et al., 2015; Nillni, 2010). Adaptive regulation of growth processes is indicated by the often-observed positive relation between ration size and growth rate in short-time experiments, e.g. in tilapia (Dong et al., 2015; Fox et al., 2010; Toguyeni et al., 1996), white sturgeon (Acipenser transmontanus) (Cui et al., 1996), gilthead sea bream (Sparus aurata) (Bermejo-Nogales et al., 2011), cod (Berg and Albert, 2003) and polar cod (Boreogadus saida) (Hop et al., 1997). Food availability is suggested to be one of the most important environmental factors influencing growth rates in fish (Dmitriew, 2011; Enberg et al., 2012; MacKenzie et al., 1998). We have not been able to find studies following hormone levels and growth rates of animals on differently sized rations throughout their growth phase.

Higher food availability in the model habitats results in higher optimal GHF levels and thus higher growth rates. Even if GHF in the model is a simplified version of the GH–IGF-1 axis, its response to stimuli like food availability resembles results from empirical studies. These studies show that concentrations of insulin-like growth factor-1 (IGF-1), a mediator of growth hormone (GH), decrease when food is less available (Bermejo-Nogales et al., 2011; Fox et al., 2010; Lescroart et al., 1998).

Even though both GH and IGF-1 are essential for growth in natural individuals, growth rate typically exhibits positive correlations with IGF-1 but not with GH (see below). In addition to promoting growth in natural fish, GH has a lipolytic effect, amplifying the use of reserves during times of food restriction (Jönsson and Björnsson, 2002). In the model, we assume stable environments and thus conflate the multiple effects of GH to a single effect on growth, thus, the lipolytic effect of GH cannot arise as a GHF-effect but would need to be prescribed through explicit assumptions.

Increasing food availability in the environment triggers high growth rates via a combined effect of THF and GHF, although THF has no direct effect on growth in the model. Empirical studies account for the effect of hormones from both hormone axes on growth, which makes the emergent correlation in THF and GHF levels plausible. Somatic growth depends on several different processes, including bone and muscle growth, which in turn combine processes regulated by hormones such as T3 and IGF-1, from the two hormone functions. A study on tilapia documented a correlation between T3 and specific growth rates (Toguyeni et al., 1996). In mammals, T3 is involved in maintenance of chondrocytes and osteoblasts (Waung et al., 2012). It may have a direct effect on bone growth by local conversion and binding to thyroid receptors or an indirect effect via GH and IGF-1 (Nilsson et al., 2005). The interplay of TH and GH is also seen in chondrocyte development, in which a first phase is triggered by IGF-1 while the second phase depends on T3 (Robson et al., 2002). The GH dynamics follow the Dual Effector Theory, in which GH can act directly on cells or indirectly via IGF-1 (Jönsson and Björnsson, 2002). Despite their actions taking place at different locations in the bones or cells, or at different times during bone maturation, bones cannot grow if one of the hormones is missing. IGF-1 also plays an important role in muscle growth (Dai et al., 2015; Grossman et al., 1997), but to our knowledge effects of thyroid on muscle growth have not been documented.

Achieving high growth rates is always related to an increased demand for energy. This demand can be met by changes in energy acquisition and allocation, and in the model we see that energy acquisition is higher in environments where food is more accessible (Fig. 3). Optimally, roughly a third of intake is allocated directly to growth while the remainders is lost to other metabolic costs on the way (Fig. 3b). The calculated average for six different teleost fish allocating metabolizable energy to growth at maximum rations of food is at about 40% (Cui and Liu, 1990). Minimum and maximum allocation rates were 21.3% and 63.4%, respectively. Thus, the optimal allocation rate found in this model is within the observed range.

From a life history perspective one would expect a decrease in length growth as the individual gets larger, due to fewer potential predators for larger fish (Bystrom et al., 2015; Persson et al., 1996) and how the increased survival prospects lead to slower optimal growth that put more weight on survival and the future. However, larger fish are more efficient feeders because they are less exposes to risk when they are foraging (Claireaux et al., 2018), countering the first effect. These two opposing forces explain the rather linear growth seen in the predicted juvenile growth from this model, an observation also seen in other adaptive models for the ontogeny of growth when acquisition is flexible (Claireaux et al., 2018; Jørgensen and Holt, 2013).

The challenges for the internal regulation mechanisms concerning storage of energy depend on the past, current, and expected food environment. In natural environments, this can include preparing for environmental change, by storing energy in reserves. In a stable food environment as in our model, building reserves is not necessary and because it involves costs it never becomes optimal, and there will be no variation in condition factor among individuals. A modelling approach analysing energy allocation in environments varying in food availability (Fischer et al., 2011) concluded that energy storage can be advantageous, but depends on the size of current reserves and how variable the environment is. An empirical study of more than 40 fish species or genera found that fish in stable habitats often have lower condition factors than fish in more unstable habitats (Fonseca and Cabral, 2007). This supports the fact that fish from the completely stable model environment have minimal reserves.

As preparation for foraging, orexin A pathways are activated when food gets scarce, while in the model impacts of OXF on intake are strongest in rich environments. In the model, we see a positive correlation between food availability and optimal OXF levels. Due to easily accessible energy in rich environments it is optimal to invest more into growth. This creates a higher energy demand in the model fish, which is met by increasing OXF levels and foraging activity. From empirical studies, orexin A is known to affect the individual’s energy budget on a short-time scale. It is negatively correlated to leptin, which serves as a proxy for the amount of stored energy in adipose tissue. Food restriction can result in higher orexin mRNA production, orexin receptor and neuron activity (Rodgers et al., 2002). This is also the case for ghrelin, acting together with orexin to prepare for and initiate foraging (Matsuda et al., 2011; Miura et al., 2007). Under fasting conditions, ghrelin levels can increase (Iwakura et al., 2015; Jönsson, 2013). Despite of the trigger, low levels of stored energy, being the same in experiments and the model, the context in which the trigger occurs is different. This results in high levels of orexin A and OXF at different food abundances.

The shift described in our model cascades from endocrinal changes affecting energy allocation and acquisition, oxygen budgets, growth, and mortality risk, which in total causes a concerted response towards more rapid growth in rich food environments. Comparing poor to rich food environments, higher growth rates are supported by THF levels that upregulate SMR and increase maximum oxygen uptake. A positive correlation between metabolic rate and a range of traits contributing to rapid growth rate was found in Trinidadian guppies *(Poecilia reticulata)* (Auer et al., 2018), and this is also the case for our model fish.

Shorter growth periods with higher growth rates in rich food environments result in higher survival. Besides supporting growth, high GHF levels contribute to reducing size-dependent mortality by growing out of vulnerable size windows more quickly. High THF levels also lower mortality, by making escapement once predators are encountered more likely to be successful. Thus, total mortality experienced through the growth phase is lower and survival at the end of the growth phase increased. To our knowledge, only GH excretion has been linked to mortality in empirical studies. The special interest assigned to GH is probably due to husbandry in which several land-living and aquatic animals have been genetically modified to excrete more GH and thus could grow faster to slaughtering size, e.g. coho salmon *(Oncorhynchus kisutch)* (Raven et al., 2008) and pig (Ju et al., 2015). Several studies have been conducted with both transgenic and hormone-implanted trout and coho salmon. Even if salmon fry can experience lower survival in the presence of predators (Sundström et al., 2005), several studies have found that fish treated with GH, thus having higher growth rates, have mortality rates similar to non-treated fish (Johnsson and Björnsson, 2001; Johnsson et al., 1999; Sundström and Devlin, 2011). In our model, these effects would come about because growth hormone increases the demand for food, and the resulting increase in appetite and foraging involves risk taking that elevates mortality rates.

The selection of fast-growing individuals over several generations also influences their endocrinology, as seen in salmon (Fleming et al., 2002). A better understanding of the combination of endocrinology and its consequences for growth is relevant also for animal breeding programs, including fish farming. Many physiological processes and traits are linked by the endocrinal network. Selecting on one of those traits will inevitably lead to changes in the endocrinal network and affect other traits. For example, selection for high growth rates could increase oxygen use in metabolic processes to a level where fish cannot sustain other metabolig processes simultaneously, something which can be described as a limited ability to multitask physiologically. This means that the majority of available oxygen is used for metabolic processes supporting growth, while little or no oxygen is left to assure free scope as is required for predator escape in the model. Other processes not modelled, like immune function, could suffer from constraints on oxygen uptake and use. A study on first-feeding salmon fry showed increases in mortality for GH-transgenic individuals under natural conditions (Sundström et al., 2004).

This model is a first step to combine internal and external control of appetite with energy allocation, growth and survival in teleost fishes. To reflect mechanisms in nature, McNamara and Houston (2009) argue that models should consist of complex environments and simplified organisms. In our case, the environment is simple while the animal model is complex. Even with this simple one-factor environment, we see a gradual change in optimal strategies for hormone expression and resulting in concerted trait differences between populations in poor and rich habitats. The model suggests an adaptive interplay of hormone functions, where GHF, OXF, and THF act together to cause an adaptive life history strategy that balances growth and survival throughout the juvenile phase. Often, effects of the internal control by means of hormones are studied in isolation from the selection pressure of the external environment. For the future, we suggest it is not sufficient to study only how hormones carry signals from tissues and sensory organs to control centres like the hypothalamus, or only how the control centre influences the decision processes in the body at many levels. Rather, there is a need to view the entire organisms as an evolved system, where key hormones mirror internal states and respond to external factors. Such decisions concern growth and survival, as in this study, but also other life history traits linked to maturation time or physiological preparations for maturation. It is this combination of emphasis on the endocrinal network in the model fish and its impacts on ultimate mechanisms as growth and survival that is characteristic of the model. It makes the model a tool for understanding processes and mechanisms underlying adaptations of growth. We think this is a fruitful path where many studies may follow.

## Acknowledgements

The authors have benefited from discussions with Sergey Budaev, Bjørn-Cato Knutsen, Tom J. Langbehn, Marc Mangel, Adèle Mennerat and Ivar Rønnestad. No competing interests declared.

## Data availability

Model code is accessible from the supplementary material, or by contacting JW.

## Appendices

**Table A1.** Parameters used in the growth model of a generalized fish using hormonal strategies to adapt to environmental challenges.

| Parameters |  |  |  |  |
| --- | --- | --- | --- | --- |
| Name | Value | Unit | Definition | Literature |
| $a$ | 0.7 | - | Exponent for standard metabolic rate | (Clarke and Johnston, 1999) |
| $b$ | 0.7 | - | Exponent for calculation of maximum aerobic scope | - |
| $d_{\text{reserves}}$ | 5 000 | $\text{J g}^{-1}$ | Energy density of reserves | - |
| $d_{\text{structure}}$ | 4 000 | $\text{J g}^{-1}$ | Energy density of soma | - |
| $k_{\text{conversion\_growth}}$ | 0.75 | - | Efficiency of converting metabolites from reserves to soma | - |
| $k_{\text{conversion\_reserves}}$ | 0.85 | - | Efficiency of converting metabolites between blood and reserves | - |
| $k_{\text{foraging}}$ | 0.2 | - | Scaling factor for energetic cost of foraging | - |
| $k_{\text{growth}}$ | 0.28 | - | Upper limit for proportional increase in structural body mass | - |
| $k_{\text{Fultons\_max}}$ | $1.2 * 10^{-8}$ | $\text{g cm}^{-1}$ | Fulton's condition factor for fish with full reserves | (Lambert and Dutil, 1997b) |
| $k_{\text{Fultons\_min}}$ | $0.85 * 10^{-8}$ | $\text{g cm}^{-1}$ | Fulton's condition factor for lean fish | (Lambert and Dutil, 1997b) |
| $k_{\text{MinutesPerWeek}}$ | 10080 | - | Number of minutes in one time step | - |
| $k_{\text{OXF}}$ | 5 | - | Scaling factor for effect of OXF on intake (including urinary and fecal energy loss) | - |
| $k_{\text{scope}}$ | $2.58 * 10^{-5}$ | $\text{J min}^{-1} \text{g}^{-b}$ | Coefficient for calculation of maximum aerobic scope | (Claireaux et al., 2000) |
| $k_{\text{SDA}}$ | 0.15 | - | Coefficient for calculation of SDA | - |
| $k_{\text{SMR}}$ | 89596.7 | $\text{J min}^{-1} \text{g}^{-a}$ | Scaling factor for standard metabolic rate | (Clarke and Johnston, 1999) |
| $k_{\text{THF\_scope}}$ | 0.24 | - | Scaling factor determining the strength of THF on AMR | - |
| $k_{\text{THF\_SMR}}$ | 0.23 | - | Scaling factor determining the strength of THF on SMR | - |
| $m_{\text{fixed}}$ | 0.0002 | $\text{year}^{-1}$ | Background mortality rate (constant) | - |
| $m_{\text{foraging}}$ | 0.08 | - | Coefficient for calculation of foraging-related mortality rate | - |
| $m_{\text{foraging} \times \text{scope}}$ | 0.9 | year | Coefficient for calculation of active-while-vulnerable mortality rate | - |
| $m_{\text{scope}}$ | 0.8 | - | Coefficient for calculation of scope-related mortality rate | - |
| $m_{\text{size}}$ | 0.038 | $\text{year}^{-1} \text{cm}^{-x_{\text{size}}}$ | Coefficient for calculation of size-dependent mortality rate | - |
| $x_{\text{foraging}}$ | 2 | - | Exponent for calculation of foraging-related mortality rate | - |
| $x_{\text{scope}}$ | 3 | - | Exponent for calculation of scope-related mortality rate | - |
| $x_{\text{size}}$ | -0.75 | - | Exponent for calculation of size-dependent mortality rate | - |
| $\alpha_{\text{max}}$ | 1500 | $\text{pg ml}^{-1}$ | Maximum value of OXF | - |
| $\gamma_{\text{max}}$ | 200 | $\text{ng ml}^{-1}$ | Maximum value of GHF | - |
| $\tau_{\text{max}}$ | 5 | $\text{ng ml}^{-1}$ | Maximum value of THF | - |

**Table A2.** Variables used in a state-dependent fish growth model using optimized hormonal strategies.

| Variables |  |  |
| --- | --- | --- |
| Name | Unit | Definition |
| $A_{\max}$ | $\text{J min}^{-1}$ | Maximum aerobic scope under influence of THF |
| $A_{\text{standard}}$ | $\text{J min}^{-1}$ | Maximum aerobic scope (AMR) |
| $B_{\text{foraging}}$ | - | Foraging behaviour |
| $C$ | $\text{J min}^{-1}$ | Energetic costs of building new tissue (soma and reserves) |
| $C_{\text{growth}}$ | J | Energy incorporated in new structural tissue |
| $E$ | - | Food abundance in environment |
| $I$ | $\text{J min}^{-1}$ | Intake (corresponds to metabolizable energy) |
| $L$ | cm | Body length |
| $M$ | $\text{year}^{-1}$ | Total mortality rate |
| $M_{\text{foraging}}$ | $\text{year}^{-1}$ | Foraging-related mortality rate |
| $M_{\text{foraging} \times \text{scope}}$ | $\text{year}^{-1}$ | Active-while-vulnerable mortality rate |
| $M_{\text{scope}}$ | $\text{year}^{-1}$ | Scope-related mortality rate |
| $M_{\text{size}}$ | $\text{year}^{-1}$ | Size-dependent mortality rate |
| $P$ | $\text{J min}^{-1}$ | Metabolic processes |
| $P_{\text{foraging}}$ | $\text{J min}^{-1}$ | Swimming cost of foraging behaviour |
| $P_{\text{growth}}$ | $\text{J min}^{-1}$ | Cost of converting metabolites from reserves into new structural tissue |
| $P_{\text{reserves}}$ | $\text{J min}^{-1}$ | Cost of converting metabolites from bloodstream into fat and proteins for storage |
| $P_{\text{SDA}}$ | $\text{J min}^{-1}$ | Cost of digestion and energy uptake into bloodstream |
| $P_{\text{SMR}}$ | $\text{J min}^{-1}$ | Standard metabolic rate (SMR) under influence of THF |
| $P_{\text{standard}}$ | $\text{J min}^{-1}$ | Standard metabolic rate (SMR) |
| $P_{\text{structure}}$ | $\text{J min}^{-1}$ | Standard metabolic rate based on structural weight |
| $R$ | J | Energy reserves |
| $R_{\max}$ | J | Maximum reserves depending on body size |
| $\Delta R$ | J | Energy incorporated in reserves (when negative, reserves are drained) |
| $S$ | $\text{year}^{-1}$ | Survival probability |
| $W$ | G | Body mass (structural and reserves) |
| $W_{\text{structure}}$ | G | Structural body mass |
| $\Delta W_{\text{structure}}$ | $\text{g week}^{-1}$ | Growth |
| $\alpha$ | $\text{pg ml}^{-1}$ | Level of OXF |
| $\gamma$ | $\text{ng ml}^{-1}$ | Level of GHF |
| $\tau$ | $\text{ng ml}^{-1}$ | Level of THF |

## References

Abbas, H. H., Authman, M. M., Zaki, M. S. and Mohamed, G. F. (2012). Effect of Seasonal Temperature Changes on Thyroid Structure and Hormones Secretion of White Grouper *(Epinephelus Aeneus)* in Suez Gulf, Egypt. Life Science Journal-Acta Zhengzhou University Overseas Edition n, 700–705.

Abbott, M. and Volkoff, H. (2011). Thyrotropin Releasing Hormone (TRH) in goldfish (*Carassius auratus*) Role in the regulation of feeding and locomotor behaviors and interactions with the orexin system and cocaine- and amphetamine regulated transcript (CART). Hormones and Behavior 59, 236–245.

Abel, E. D., Ahima, R. S., Boers, M. E., Elmquist, J. K. and Wondisford, F. E. (2001). Critical role for thyroid hormone receptor beta 2 in the regulation of paraventricular thyrotropin-releasing hormone neurons. Journal of Clinical Investigation 107,1017–1023.

Ali, M., Nicieza, A. and Wootton, R. J. (2003). Compensatory growth in fishes: a response to growth depression. Fish and Fisheries 4,147–190.

Andersen, B. S., Jørgensen, C., Eliassen, S. and Giske, J. (2016). The proximate architecture for decision-making in fish. Fish and Fisheries 17, 680–695.

Auer, S. K., Dick, C. A., Metcalfe, N. B. and Reznick, D. N. (2018). Metabolic rate evolves rapidly and in parallel with the pace of life history. Nature Communications 9, 6.

Berg, E. and Albert, O. T. (2003). Cod in fjords and coastal waters of North Norway: distribution and variation in length and maturity at age. Ices Journal of Marine Science 60, 787–797.

Bermejo-Nogales, A., Benedito-Palos, L., Calduch-Giner, J. A. and Perez-Sanchez, J. (2011). Feed restriction up-regulates uncoupling protein 3 (UCP3) gene expression in heart and red muscle tissues of gilthead sea bream (*Sparus aurata* L.) New insights in substrate oxidation and energy expenditure. Comparative Biochemistry and Physiology a-Molecular & Integrative Physiology 159, 296–302.

Björnsson, B. T. (1997). The biology of salmon growth hormone: from daylight to dominance. Fish Physiology and Biochemistry 17, 9–24.

Brown, J. H., Gillooly, J. F., Allen, A. P., Savage, V. M. and West, G. B. (2004). Toward a metabolic theory of ecology. Ecology 85,1771–1789.

Bubenik, G. A., Schams, D., White, R. G., Rowell, J., Blake, J. and Bartos, L. (1998). Seasonal levels of metabolic hormones and substrates in male and female reindeer (*Rangifer taranduś)*. Comparative Biochemistry and Physiology C-Pharmacology Toxicology & Endocrinology 120, 307–315.

Budaev, S., Jørgensen, C., Mangel, M., Eliassen, S. and Giske, J. (2019). Decision-Making From the Animal Perspective: Bridging Ecology and Subjective Cognition. Frontiers in Ecology and Evolution 7.

Bureau, D. P., Kaushik, S. J. and Cho, C. Y. (2003). 1 - Bioenergetics A2 - Halver, John E. In Fish Nutrition (Third Edition), (ed. R. W. Hardy), pp. 1–59. San Diego: Academic Press.

Burrows, M. T. (1994). An optimal foraging and migration model for juvenile plaice. Evolutionary Ecology 8,125–149.

Bystrom, P., Bergstrom, U., Hjalten, A., Stahl, S., Jonsson, D. and Olsson, J. (2015). Declining coastal piscivore populations in the Baltic Sea: Where and when do sticklebacks matter? Ambio 44, S462–S471.

Cabello, G. and Wrutniak, C. (1989). Thyroid-hormone and growth: relationships with growth hormone effects and regulation. Reproduction Nutrition Development 29, 387–402.

Cai, X. J., Evans, M. L., Lister, C. A., Leslie, R. A., Arch, J. R. S., Wilson, S. and Williams, G. (2001). Hypoglycemia activates orexin neurons and selectively increases hypothalamic orexin-B levels - Responses inhibited by feeding and possibly mediated by the nucleus of the solitary tract. Diabetes 50,105–112.

Cai, X. J., Liu, X. H., Evans, M., Clapham, J. C., Wilson, S., Arch, J. R. S., Morris, R. and Williams, G. (2002). Orexins and feeding: special occasions or everyday occurrence? Regulatory Peptides 104,1–9.

Cai, X. J., Widdowson, P. S., Harrold, J., Wilson, S., Buckingham, R. E., Arch, J. R. S., Tadayyon, M., Clapham, J. C., Wilding, J. and Williams, G. (1999). Hypothalamic orexin expression - Modulation by blood glucose and feeding. Diabetes 48, 2132–2137.

Cameron, C., Moccia, R., Azevedo, P. A. and Leatherland, J. F. (2007). Effect of diet and ration on the relationship between plasma GH and IGF-1 concentrations in Arctic charr, *Salvelinus alpinus* (L.). Aquaculture Research 38,877–886.

Cammisotto, P. G. and Bendayan, M. (2007). Leptin secretion by white adipose tissue and gastric mucosa. Histology and Histopathology 22,199–210.

Carr, A. N. and Kranias, E. G. (2002). Thyroid hormone regulation of calcium cycling proteins. Thyroid 12, 453–457.

Chatterjee, A., Hsieh, Y. L. and Yu, J. Y. L. (2001). Molecular cloning of cDNA encoding thyroid stimulating hormone beta subunit of bighead carp *Aristichthys nobilis* and regulation of its gene expression. Molecular and Cellular Endocrinology 174,1–9.

Claireaux, G., Webber, D. M., Lagardere, J. P. and Kerr, S. R. (2000). Influence of water temperature and oxygenation on the aerobic metabolic scope of Atlantic cod (*Gadus morhua)*. Journal of Sea Research 44, 257–265.

Claireaux, M., Jørgensen, C. and Enberg, K. (2018). Evolutionary effects of fishing gear on foraging behavior and life-history traits. Ecology and Evolution 8,10711–10721.

Clark, C. W. and Mangel, M. (2000). Dynamic State Variable Models in Ecology. Cary: Oxford University Press, USA.

Clarke, A. and Johnston, N. M. (1999). Scaling of metabolic rate with body mass and temperature in teleost fish. Journal of Animal Ecology 68, 893–905.

Cui, Y., Hung, S. S. O. and Zhu, X. (1996). Effect of ration and body size on the energy budget of juvenile white sturgeon. Journal of Fish Biology 49, 863–876.

Cui, Y. and Liu, J. K. (1990). Comparison of energy budget among six teleosts - III. Growth rate and energy budget. Comparative Biochemistry and Physiology a-Physiology 97, 381–384.

Dai, X. Y., Zhang, W., Zhuo, Z. J., He, J. Y. and Yin, Z. (2015). Neuroendocrine regulation of somatic growth in fishes. Science China-Life Sciences 58,137–147.

Dennett, D. C. (2017). From bacteria to Bach and back: the evolution of minds. London: Allen Lane.

Di Liegro, I. (2008). Thyroid hormones and the central nervous system of mammals (Review). Molecular Medicine Reports 1, 279–295.

Dmitriew, C. M. (2011). The evolution of growth trajectories: what limits growth rate? Biological Reviews 86, 97–116.

Dong, G. F., Yang, Y. O., Yao, F., Chen, L., Bu, F. Y., Li, P. C., Huang, F. and Yu, D. H. (2015). Individual variations and interrelationships in feeding rate, growth rate, and spontaneous activity in hybrid tilapia (*Oreochromis niloticus x 0. aureus*) at different feeding frequencies. Journal of Applied Ichthyology 31, 349–354.

Dube, M. G., Kalra, S. P. and Kalra, P. S. (1999). Food intake elicited by central administration of orexins/hypocretins: identification of hypothalamic sites of action. Brain Research 842, 473–477.

Eales, J. G. (1988). The Influence of Nutritional State on Thyroid Function in Various Vertebrates. American Zoologist 28, 351–362.

Edwards, C. M. B., Abusnana, S., Sunter, D., Murphy, K. G., Ghatel, M. A. and Bloom, S. R. (1999). The effect of the orexins on food intake: comparison with neuropeptide Y, melanin-concentrating hormone and galanin. Journal of Endocrinology 160, R7–R12.

Elton, C. S. (1927). Animal ecology. London: Sidgwick & Jackson.

Enberg, K., Jørgensen, C., Dunlop, E. S., Varpe, Ø., Boukal, D. S., Baulier, L., Eliassen, S. and Heino, M. (2012). Fishing-induced evolution of growth: concepts, mechanisms and the empirical evidence. Marine Ecology-an Evolutionary Perspective 33,1–25.

Facciolo, R. M., Crudo, M., Giusi, G. and Canonaco, M. (2010). GABAergic influences on ORX receptor-dependent abnormal motor behaviors and neurodegenerative events in fish. Toxicology and Applied Pharmacology 243, 77–86.

Fawcett, T. W., Fallenstein, B., Higginson, A. D., Houston, A. I., Mallpress, D. E. W., Trimmer, P. C. and McNamara, J. M. (2014). The evolution of decision rules in complex environments. Trends in Cognitive Sciences 18,153–161.

Fekete, C. and Lechan, R. M. (2014). Central Regulation of Hypothalamic-Pituitary-Thyroid Axis Under Physiological and Pathophysiological Conditions. Endocrine Reviews 35,159–194.

Fiksen, Ø., Giske, J. and Slagstad, D. (1995). A spatially explicit fitness based model of capelin migrations in the barents sea. Fisheries Oceanography 4,193–208.

Fischer, B., Dieckmann, U. and Taborsky, B. (2011). WHEN TO STORE ENERGY IN A STOCHASTIC ENVIRONMENT. Evolution 65,1221–1232.

Fisher, R. A. (1930). Genetical theory of natural selection Oxford, U: Oxford University Press.

Fleming, I. A., Agustsson, T., Finstad, B., Johnsson, J. I. and Björnsson, B. T. (2002). Effects of domestication on growth physiology and endocrinology of Atlantic salmon (Salmo salar). Canadian Journal of Fisheries and Aquatic Sciences 59,1323–1330.

Fonseca, V. F. and Cabral, H. N. (2007). Are fish early growth and condition patterns related to life-history strategies? Reviews in Fish Biology and Fisheries 17, 545–564.

Fox, B. K., Breves, J. P., Davis, L. K., Pierce, A. L., Hirano, T. and Grau, E. G. (2010). Tissue-specific regulation of the growth hormone/insulin-like growth factor axis during fasting and re-feeding: Importance of muscle expression of IGF-I and IGF-II mRNA in the tilapia. General and Comparative Endocrinology 166, 573–580.

Friesema, E. C. H., Docter, R., Moerings, E., Stieger, B., Hagenbuch, B., Meier, P. J., Krenning, E. P., Hennemann, G. and Visser, T. J. (1999). Identification of thyroid hormone transporters. Biochemical and Biophysical Research Communications 254, 497–501.

Fry, F. E. J. (1971). The effect of environmental factors on the physiology of fish. In Fish Physiology vol. 6 eds. W. S. Hoar and D. J. Randall), pp. 1–98: Academic Press.

Gahete, M. D., Duran-Prado, M., Luque, R. M., Martinez-Fuentes, A. J., Quintero, A., Gutierrez-Pascual, E., Cordoba-Chacon, J., Malagon, M. M., Gracia-Navarro, F. and Castano, J. P. (2009). Understanding the Multifactorial Control of Growth Hormone Release by Somatotropes Lessons from Comparative Endocrinology. In Trends in Comparative Endocrinology and Neurobiology, vol. 1163 eds. H. Vaudry E. W. Roubos G. M. Coast and M. Vallarino), pp. 137–153. Malden: Wiley-Blackwell.

Garland, T., Zhao, M. and Saltzman, W. (2016). Hormones and the Evolution of Complex Traits: Insights from Artificial Selection on Behavior. Integrative and Comparative Biology 56, 207–224.

Gatford, K. L., Egan, A. R., Clarke, I. J. and Owens, P. C. (1998). Sexual dimorphism of the somatotrophic axis. Journal of Endocrinology 157, 373–389.

Giske, J., Eliassen, S., Fiksen, O., Jakobsen, P. J., Aksnes, D. L., Jorgensen, C. and Mangel, M. (2013). Effects of the Emotion System on Adaptive Behavior. American Naturalist 182, 689–703.

Gislason, H., Daan, N., Rice, J. C. and Pope, J. G. (2010). Size, growth, temperature and the natural mortality of marine fish. Fish and Fisheries 11, 149–158.

Grafen, A. (1984). Natural selection, kin selection and group selection, eds. K. Jr and D. Nb), pp. 62–84. Oxford: Blackwell Scientific Publications.

Grossman, E. J., Grindeland, R. E., Roy, R. R., Talmadge, R. J., Evans, J. and Edgerton, V. R. (1997). Growth hormone, IGF-I, and exercise effects on non-weight-bearing fast muscles of hypophysectomized rats. Journal of Applied Physiology 83,1522–1530.

Hackney, A. C. and Dobridge, J. D. (2009). Thyroid hormones and the interrelationship of cortisol and prolactin: influence of prolonged, exhaustive exercise. Endokrynologia Polska 60, 252–257.

Hamrick, M. W. and Ferrari, S. L. (2008). Leptin and the sympathetic connection of fat to bone. Osteoporosis International 19, 905–912.

Han, Y. S., Liao, I. C., Tzeng, W. N. and Yu, J. Y. L. (2004). Cloning of the cDNA for thyroid stimulating hormone beta subunit and changes in activity of the pituitary-thyroid axis during silvering of the Japanese eel, *Anguilla japonica*. Journal of Molecular Endocrinology 32,179–194.

Haynes, A. C., Jackson, B., Overend, P., Buckingham, R. E., Wilson, S., Tadayyon, M. and Arch, J. R. S. (1999). Effects of single and chronic intracerebroventricular administration of the orexins on feeding in the rat. Peptides 20,1099–1105.

Heilbronn, L. K., de Jonge, L., Frisard, M. I. and et al. (2006). Effect of 6-month calorie restriction on biomarkers of longevity, metabolic adaptation, and oxidative stress in overweight individuals: A randomized controlled trial. JAMA 295,1539–1548.

Herwig, A., Ross, A. W., Nilaweera, K. N., Morgan, P. J. and Barrett, P. (2008). Hypothalamic Thyroid Hormone in Energy Balance Regulation. Obesity Facts 1, 71–79.

Holdway, D. A. and Beamish, F. W. H. (1984). Specific growth rate and proximate body composition of atlantic cod (*Gadus morhua* L). Journal of Experimental Marine Biology and Ecology 81,147–170.

Holt, R. E. and Jørgensen, C. (2014). Climate warming causes life-history evolution in a model for Atlantic cod (Gadus morhua). Conservation Physiology 2.

Holt, R. E. and Jørgensen, C. (2015). Climate change in fish: effects of respiratory constraints on optimal life history and behaviour. Biology Letters 11.

Hop, H., Tonn, W. M. and Welch, H. E. (1997). Bioenergetics of Arctic cod *(Boreogadus saida)* at low temperatures. Canadian Journal of Fisheries and Aquatic Sciences 54,1772–1784.

Horvath, T. L., Diano, S. and van den Pol, A. N. (1999). Synaptic interaction between hypocretin (Orexin) and neuropeptide Y cells in the rodent and primate hypothalamus: A novel circuit implicated in metabolic and endocrine regulations. Journal of Neuroscience 19,1072–1087.

Hoskins, L. J., Xu, M. Y. and Volkoff, H. (2008). Interactions between gonadotropin-releasing hormone (GnRH) and orexin in the regulation of feeding and reproduction in goldfish (*Carassius auratus)*. Hormones and Behavior 54, 379–385.

Houston, A. I. and McNamara, J. M. (1999). Models of Adaptive Behaviour: An Approach Based on State. Cambridge, UK: Cambridge University Press.

Ikegami, K. and Yoshimura, T. (2013). Seasonal Time Measurement During Reproduction. The Journal of Reproduction and Development 59, 327–333.

Iwakura, H., Kangawa, K. and Nakao, K. (2015). The regulation of circulating ghrelin - with recent updates from cell-based assays. Endocrine Journal 62,107–122.

Jenni-Eiermann, S., Jenni, L. and Piersma, T. (2002). Temporal uncoupling of thyroid hormones in Red Knots: T3 peaks in cold weather, T4 during moult. Journal Fur Ornithologie 143, 331–340.

Johnsson, J. I. and Björnsson, B. T. (2001). Growth-enhanced fish can be competitive in the wild. Functional Ecology 15, 654–659.

Johnsson, J. I., Petersson, E., Jönsson, E., Järvi, T. and Björnsson, B. T. (1999). Growth hormone-induced effects on mortality, energy status and growth: a field study on Brown Trout (*Salmo trutta)*. Functional Ecology 13, 514–522.

Johnston, I. A. (1981). Structure and function of fish muscle. Symp Zool Soc Lond 48, 71–113.

Ju, H. M., Zhang, J. Q., Bai, L. J., Mu, Y. L., Du, Y. T., Yang, W. X., Li, Y., Sheng, A. Z. and Li, K. (2015). The transgenic cloned pig population with integrated and controllable GH expression that has higher feed efficiency and meat production. Scientific Reports 5,14.

Jönsson, E. (2013). The role of ghrelin in energy balance regulation in fish. General and Comparative Endocrinology 187, 79–85.

Jönsson, E. and Björnsson, B. T. (2002). Physiological functions of growth hormone in fish with special reference to its influence on behaviour. Fisheries Science 68, 742–748.

Jørgensen, C. and Holt, R. E. (2013). Natural mortality: Its ecology, how it shapes fish life histories, and why it may be increased by fishing. Journal of Sea Research 75, 8–18.

Jørgensen, E. H. and Johnsen, H. K. (2014). Rhythmic life of the Arctic charr: Adaptations to life at the edge. Marine Genomics 14, 71–81.

Kalamatianos, T., Markianos, M., Margetis, K., Bourlogiannis, F. and Stranjalis, G. (2014). Higher Orexin A levels in lumbar compared to ventricular CSF: A study in idiopathic normal pressure hydrocephalus. Peptides 51,1–3.

Kalra, S. P., Dube, M. G., Pu, S. Y., Xu, B., Horvath, T. L. and Kalra, P. S. (1999). Interacting appetite-regulating pathways in the hypothalamic regulation of body weight. Endocrine Reviews 20, 68–100.

Kaminski, T., Nitkiewicz, A. and Smolinska, N. (2013). Changes in plasma orexin A and orexin B concentrations during the estrous cycle of the pig. Peptides 39,175–177.

Killen, S. S., Costa, I., Brown, J. A. and Gamperl, A. K. (2007). Little left in the tank: metabolic scaling in marine teleosts and its implications for aerobic scope. Proceedings of the Royal Society B-Biological Sciences 274, 431–438.

Killen, S. S., Mitchell, M. D., Rummer, J. L., Chivers, D. P., Ferrari, M. C. O., Meekan, M. G. and McCormick, M. I. (2014). Aerobic scope predicts dominance during early life in a tropical damselfish. Functional Ecology 28, 1367–1376.

Kirby, D. S., Fiksen, O. and Hart, P. J. B. (2000). A dynamic optimisation model for the behaviour of tunas at ocean fronts. Fisheries Oceanography 9, 328–342.

Kitano, J., Lema, S. C., Luckenbach, J. A., Mori, S., Kawagishi, Y., Kusakabe, M., Swanson, P. and Peichel, C. L. (2010). Adaptive Divergence in the Thyroid Hormone Signaling Pathway in the Stickleback Radiation. Current Biology 20, 2124–2130.

Kjesbu, O. S., Klungsøyr, J., Kryvi, H., Witthames, P. R. and Walker, M. G. (1991). Fecundity, atresia, and egg size of captive Atlantic cod (*Gadus morhua*) in relation to proximate body composition. Canadian Journal of Fisheries and Aquatic Sciences 48, 2333–2343.

Kojima, M., Hosoda, H., Date, Y., Nakazato, M., Matsuo, H. and Kangawa, K. (1999). Ghrelin is a growth-hormone-releasing acylated peptide from stomach. Nature 402, 656–660.

Kooijman, S. (2001). Quantitative aspects of metabolic organization: a discussion of concepts. Philosophical Transactions of the Royal Society of London Series B-Biological Sciences 356, 331–349.

Kooijman, S. A. L. M. (1993). Dynamic energy budgets in biological systems: theory and applications in ecotoxicology. Cambridge: Cambridge University Press.

Krause, J. and Godin, J. G. J. (1996). Influence of prey foraging posture on flight behavior and predation risk: Predators take advantage of unwary prey. Behavioral Ecology 7, 264–271.

Kuba, J., Blaszczyk, B., Stankiewicz, T., Skuratko, A. and Udala, J. (2015). Analysis of annual changes in the concentrations of selected macro- and microelements, thyroxine, and testosterone in the serum of Red Deer (*Cervus elaphus*) stags. Biological Trace Element Research 168, 356–361.

Lambert, Y. and Dutil, J. D. (1997a). Can simple condition indices be used to monitor and quantify seasonal changes in the energy reserves of cod (Gadus morhua)? Canadian Journal of Fisheries and Aquatic Sciences 54, 104–112.

Lambert, Y. and Dutil, J. D. (1997b). Condition and energy reserves of Atlantic cod (*Gadus morhua*) during the collapse of the northern Gulf of St. Lawrence stock. Canadian Journal of Fisheries and Aquatic Sciences 54, 2388–2400.

Lanfranco, F., Gianotti, L., Giordano, R., Pellegrino, M., Maccario, M. and Arvat, E. (2003). Ageing, growth hormone and physical performance. Journal of Endocrinological Investigation 26, 861–872.

Lanni, A., Moreno, M., Lombardi, A., de Lange, P. and Goglia, F. (2001). Control of energy metabolism by iodothyronines. Journal of Endocrinological Investigation 24, 897–913.

Lartey, L. J., Werneck-de-Castro, J. P., O-Sullivan, I., Unterman, T. G. and Bianco, A. C. (2015). Coupling between Nutrient Availability and Thyroid Hormone Activation. Journal of Biological Chemistry 290, 30551–30561.

Lescroart, O., Roelants, I., Darras, V. M., Kuhn, E. R. and Ollevier, F. (1998). Apomorphine-induced growth hormone release in African catfish (*Clarias gariepinus*) is related to the animal’s nutritional status. Fish Physiology and Biochemistry 18, 353–361.

Lima, S. L. (1998). Stress and decision making under the risk of predation: Recent developments from behavioral, reproductive, and ecological perspectives. Stress and Behavior 27, 215–290.

Lima, S. L. and Dill, L. M. (1990). Behavioral decisions made under the risk of predation: a review and prospectus. Canadian Journal of Zoology-Revue Canadienne De Zoologie 68, 619–640.

Little, A. G. and Seebacher, F. (2014). Thyroid hormone regulates cardiac performance during cold acclimation in zebrafish (*Danio rerio)*. Journal of Experimental Biology 217, 718–725.

Lubkin, M. and Stricker-Krongrad, A. (1998). Independent feeding and metabolic actions of orexins in mice. Biochemical and Biophysical Research Communications 253, 241–245.

MacKenzie, D. S., VanPutte, C. M. and Leiner, K. A. (1998). Nutrient regulation of endocrine function in fish. Aquaculture 161, 3–25.

Malik, R. and Hodgson, H. (2002). The relationship between the thyroid gland and the liver. Qjm-an International Journal of Medicine 95, 559–569.

Marras, S., Claireaux, G., McKenzie, D. J. and Nelson, J. A. (2010). Individual variation and repeatability in aerobic and anaerobic swimming performance of European sea bass, Dicentrarchus labrax. Journal of Experimental Biology 213, 26–32.

Matsuda, K., Azuma, M. and Kang, K. S. (2012). Orexin system in teleost fish. In Vitamins and Hormones: Sleep Hormones, Vol 89, vol. 89 (ed. G. Litwack), pp. 341–361. San Diego: Elsevier Academic Press Inc.

Matsuda, K., Kang, K. S., Sakashita, A., Yahashi, S. and Vaudry, H. (2011). Behavioral effect of neuropeptides related to feeding regulation in fish. In Trends in Neuroendocrinology, vol. 1220 eds. H. Vaudry and M. C. Tonon), pp. 117–126. Oxford: Blackwell Science Publ.

McNamara, J. M. and Houston, A. I. (2009). Integrating function and mechanism. Trends in Ecology & Evolution 24, 670–675.

Miura, M., Tanaka, K., Komatsu, Y., Suda, M., Yasoda, A., Sakuma, Y., Ozasa, A. and Nakao, K. (2002). Thyroid hormones promote chondrocyte differentiation in mouse ATDC5 cells and stimulate endochondral ossification in fetal mouse tibias through lodothyronine deiodinases in the growth plate. Journal of Bone and Mineral Research 17, 443–454.

Miura, T., Maruyama, K., Shimakura, S. I., Kaiya, H., Uchiyama, M., Kangawa, K., Shioda, S. and Matsuda, K. (2007). Regulation of food intake in the goldfish by interaction between ghrelin and orexin. Peptides 28, 1207–1213.

Mozsar, A., Boros, G., Saly, P., Antal, L. and Nagy, S. A. (2015). Relationship between Fulton’s condition factor and proximate body composition in three freshwater fish species. Journal of Applied Ichthyology 31,315–320.

Mullur, R., Liu, Y.-Y. and Brent, G. A. (2014). Thyroid Hormone Regulation of Metabolism. Physiological Reviews 94, 355–382.

Müller, T. D., Nogueiras, R., Andermann, M. L., Andrews, Z. B., Anker, S. D., Argente, J., Batterham, R. L., Benoit, S. C., Bowers, C. Y., Broglio, F. et al. (2015). Ghrelin. Molecular Metabolism 4, 437–460.

Nakamachi, T., Matsuda, K., Maruyama, K., Miura, T., Uchiyama, M., Funahashi, H., Sakurai, T. and Shioda, S. (2006). Regulation by orexin of feeding behaviour and locomotor activity in the goldfish. Journal of Neuroendocrinology 18, 290–297.

Nillni, E. A. (2010). Regulation of the hypothalamic Thyrotropin Releasing Hormone (TRH) neuron by neuronal and peripheral inputs. Frontiers in Neuroendocrinology 31,134–156.

Nillni, E. A., Vaslet, C., Harris, M., Hollenberg, A., Bjørbæk, C. and Flier, J. S. (2000). Leptin regulates prothyrotropin-releasing hormone biosynthesis - Evidence for direct and indirect pathways. Journal of Biological Chemistry 275, 36124–36133.

Nilsson, O., Marino, R., De Luca, F., Phillip, M. and Baron, J. (2005). Endocrine regulation of the growth plate. Hormone Research 64,157–165.

Nisbet, R. M., Muller, E. B., Lika, K. and Kooijman, S. (2000). From molecules to ecosystems through dynamic energy budget models. Journal of Animal Ecology 69, 913–926.

Oka, Y., Kanbayashi, T., Mezaki, T., Iseki, K., Matsubayashi, J., Murakami, G., Matsui, M., Shimizu, T. and Shibasaki, H. (2004). Low CSF hypocretin-1/orexin-A associated with hypersomnia secondary to hypothalamic lesion in a case of multiple sclerosis. Journal of Neurology 251,885–886.

Ono, H., Hoshino, Y., Yasuo, S., Watanabe, M., Nakane, Y., Murai, A., Ebihara, S., Korf, H. W. and Yoshimura, T. (2008). Involvement of thyrotropin in photoperiodic signal transduction in mice. Proceedings of the National Academy of Sciences of the United States of America 105,18238–18242.

Pangle, K. L. and Sutton, T. M. (2005). Temporal changes in the relationship between condition indices and proximate composition of juvenile *Coregonus artedi*. Journal of Fish Biology 66,1060–1072.

Penney, C. C. and Volkoff, H. (2014). Peripheral injections of cholecystokinin, apelin, ghrelin and orexin in cavefish (Astyanax fasciatus mexicanus): Effects on feeding and on the brain expression levels of tyrosine hydroxylase, mechanistic target of rapamycin and appetite-related hormones. General and Comparative Endocrinology 196, 34–40.

Persson, L., Andersson, J., Wahlstrom, E. and Eklov, P. (1996). Size-specific interactions in lake systems: Predator gape limitation and prey growth rate and mortality. Ecology 77, 900–911.

Peter, R. E. and Marchant, T. A. (1995). The endocrinology of growth in carp and related species. Aquaculture 129, 299–321.

Peterson, I. and Wroblewski, J. S. (1984). Mortality rate of fishes in the pelagic ecosystem. Canadian Journal of Fisheries and Aquatic Sciences 41, 1117–1120.

Power, D. M., Melo, J. and Santos, C. R. A. (2000). The effect of food deprivation and refeeding on the liver, thyroid hormones and transthyretin in sea bream. Journal of Fish Biology 56, 374–387.

Priede, I. G. (1985). Metabolic scope in fishes. In Fish energetics: New perspectives, eds. P. Tytler and P. Calow), pp. 33–64. Baltimor, MA: John Hopkins University Press

Raven, P. A., Uh, M., Sakhrani, D., Beckman, B. R., Cooper, K., Pinter, J., Leder, E. H., Silverstein, J. and Devlin, R. H. (2008). Endocrine effects of growth hormone overexpression in transgenic coho salmon. General and Comparative Endocrinology 159, 26–37.

Reale, D., Garant, D., Humphries, M. M., Bergeron, P., Careau, V. and Montiglio, P. O. (2010). Personality and the emergence of the pace-of-life syndrome concept at the population level. Philosophical Transactions of the Royal Society B-Biological Sciences 365, 4051–4063.

Reznick, D., Nunney, L. and Tessier, A. (2000). Big houses, big cars, superfleas and the costs of reproduction. Trends in Ecology & Evolution 15, 421–425.

Richmond, J. P., Du Dot, T. J., Rosen, D. A. S. and Zinn, S. A. (2010). Seasonal Influence on the Response of the Somatotropic Axis to Nutrient Restriction and Re-alimentation in Captive Steller Sea Lions (*Eumetopias jubatuś)*. Journal of Experimental Zoology Part a-Ecological Genetics and Physiology 313A, 144–156.

Robson, H., Siebler, T., Shalet, S. M. and Williams, G. R. (2002). Interactions between GH, IGF-I, glucocorticoids, and thyroid hormones during skeletal growth. Pediatric Research 52.

Rodgers, R. J., Halford, J. C. G., de Souza, R. L. N., de Souza, A. L. C., Piper, D. C., Arch, J. R. S. and Blundell, J. E. (2000). Dose-response effects of orexin-A on food intake and the bahavioural satiety sequence in rats. Regulatory Peptides 96, 71–84.

Rodgers, R. J., Ishii, Y., Halford, J. C. G. and Blundell, J. E. (2002). Orexins and appetite regulation. Neuropeptides 36, 303–325.

Rome, L. C., Funke, R. P., Alexander, R. M., Lutz, G., Aldridge, H., Scott, F. and Freadman, M. (1988). WHY ANIMALS HAVE DIFFERENT MUSCLE-FIBER TYPES. Nature 335, 824–827.

Rountree, R. A. and Sedberry, G. R. (2009). A theoretical model of shoaling behavior based on a consideration of patterns of overlap among the visual fields of individual members. Acta Ethologica 12, 61–70.

Rønnestad, I., Gomes, A. S., Murashita, K., Angotzi, R., Jonsson, E. and Volkoff, H. (2017). Appetite-Controlling Endocrine Systems in Teleosts. Frontiers in Endocrinology 8, 24.

Sakurai, T., Amemiya, A., Ishii, M., Matsuzaki, I., Chemelli, R. M., Tanaka, H., Williams, S. C., Richardson, J. A., Kozlowski, G. P., Wilson, S. et al. (1998). Orexins and orexin receptors: A family of hypothalamic neuropeptides and G protein-coupled receptors that regulate feeding behavior. Cell 92, 573–585.

Salzman, T. C., McLaughlin, A. L., Westneat, D. F. and Crowley, P. H. (2018). Energetic trade-offs and feedbacks between behavior and metabolism influence correlations between pace-of-life attributes. Behavioral Ecology and Sociobiology 72,18.

Sih, A. (1992). PREY UNCERTAINTY AND THE BALANCING OF ANTIPREDATOR AND FEEDING NEEDS. American Naturalist 139,1052–1069.

Silberman, D. M., Wald, M. and Genaro, A. M. (2002). Effects of chronic mild stress on lymphocyte proliferative response. Participation of serum thyroid hormones and corticosterone. International Immunopharmacology 2,487–497.

Silva, J. E. (2003). The Thermogenic Effect of Thyroid Hormone and Its Clinical Implications. Annals of Internal Medicine 139, 205–213.

Sinervo, B. and Svensson, E. (1998). Mechanistic and selective causes of life history trade-offs and plasticity. Oikos 83, 432–442.

Sundström, L. F. and Devlin, R. H. (2011). Increased intrinsic growth rate is advantageous even under ecologically stressful conditions in coho salmon (*Oncorhynchus kisutch)*. Evolutionary Ecology 25, 447–460.

Sundström, L. F., Lohmus, M. and Devlin, R. H. (2005). Selection on increased intrinsic growth rates in coho salmon, *Oncorhynchus kisutch*. Evolution 59,1560–1569.

Sundström, L. F., Lohmus, M., Johnsson, J. I. and Devlin, R. H. (2004). Growth hormone transgenic salmon pay for growth potential with increased predation mortality. Proceedings of the Royal Society B-Biological Sciences 271, S350–S352.

Suttie, J. M., Breier, B. H., Gluckman, P. D., Littlejohn, R. P. and Webster, J. R. (1992). Effects of melatonin implants on insulin-like growth factor 1 in male red deer (*Cervus elaphuś)*. General and Comparative Endocrinology 87,111–119.

Suttie, J. M., White, R. G., Breier, B. H. and Gluckman, P. D. (1991). Photoperiod associated changes in insulin-like growth factor-I in reindeer. Endocrinology 129, 679–682.

Suttie, J. M., White, R. G., Manley, T. R., Breier, B. H., Gluckman, P. D., Fennessy, P. F. and Woodford, K. (1993). Insulin-like growth factor 1 and growth seasonality in reindeer (*Rangifer tarandus*) - comparisons with temperate and tropical cervids. Rangifer 13.

Tagawa, M., Ogasawara, T., Sakamoto, T., Miura, T., Yamauchi, K. and Hirano, T. (1994). Thyroid hormone concentrations in the gonads of wild chum salmon during maturation. Fish Physiology and Biochemistry 13, 233–240.

Tinbergen, N. (1963). On Aims and Methods of Ethology. Zeitschrift für Tierpşychologie 20, 410–433.

Toguyeni, A., Baroiller, J. F., Fostier, A., LeBail, P. Y., Kuhn, E. R., Mol, K. A. and Fauconneau, B. (1996). Consequences of food restriction on short-term growth variation and on plasma circulating hormones in *Oreochromis niloticus* in relation to sex. General and Comparative Endocrinology 103,167–175.

Tomasik, P. J., Spodaryk, M. and Sztefko, K. (2004). Plasma concentrations of orexins in children. Annals of Nutrition and Metabolism 48,215–220.

Uchida, K., Kajimura, S., Riley, L. G., Hirano, T., Aida, K. and Grau, E. G. (2003). Effects of fasting on growth hormone/insulin-like growth factor I axis in the tilapia, *Oreochromis mossambicus*. Comparative Biochemistry and Physiology Part A: Molecular & Integrative Physiology 134, 429–439.

Van der Geyten, S., Mol, K. A., Pluymers, W., Kuhn, E. R. and Darras, V. M. (1998). Changes in plasma T-3 during fasting/refeeding in tilapia *[Oreochromis niloticus*) are mainly regulated through changes in hepatic type II iodothyronine deiodinase. Fish Physiology and Biochemistry 19,135–143.

van der Post, D. J. and Semmann, D. (2011). Patch depletion, niche structuring and the evolution of co-operative foraging. Bmc Evolutionary Biology 11,16.

Varghese, S. and Oommen, O. V. (1999). Thyroid hormones regulate lipid metabolism in a teleost *Anabas testudineus* (Bloch). Comparative Biochemistry and Physiology B-Biochemistry & Molecular Biology 124, 445–450.

Varghese, S., Shameena, B. and Oommen, O. V. (2001). Thyroid hormones regulate lipid peroxidation and antioxidant enzyme activities in *Anabas testudineus* (Bloch). Comparative Biochemistry and Physiology B-Biochemistry & Molecular Biology 128,165–171.

Velez, E. J., Perello-Amoros, M., Lutfi, E., Azizi, S., Capilla, E., Navarro, I., Perez-Sanchez, J., Calduch-Giner, J. A., Blasco, J., Fernandez-Borras, J. et al. (2019). A long-term growth hormone treatment stimulates growth and lipolysis in gilthead sea bream juveniles. Comparative Biochemistry and Physiology a-Molecular & Integrative Physiology 232, 67–78.

Vermeulen, A. (2002). Ageing, hormones, body composition, metabolic effects. World Journal of Urology 20, 23–27.

Volkoff, H., Bjorklund, J. M. and Peter, R. E. (1999). Stimulation of feeding behavior and food consumption in the goldfish, *Carassius auratus,* by orexin-A and orexin-B. Brain Research 846, 204–209.

Volkoff, H., Eykelbosh, A. J. and Peter, R. E. (2003). Role of leptin in the control of feeding of goldfish *Carassius auratus·.* interactions with cholecystokinin, neuropeptide Y and orexin A, and modulation by fasting. Brain Research 972, 90–109.

Waung, J. A., Bassett, J. H. D. and Williams, G. R. (2012). Thyroid hormone metabolism in skeletal development and adult bone maintenance. Trends in Endocrinology and Metabolism 23,155–162.

Webb, P. (2004). Selective activators of thyroid hormone receptors. Expert Opinion on Investigational Drugs 13, 489–500.

Weber, J. M., Choi, K., Gonzalez, A. and Omlin, T. (2016). Metabolic fuel kinetics in fish: swimming, hypoxia and muscle membranes. Journal of Experimental Biology 219, 250–258.

Yang, B. Y., Zhai, G., Gong, Y. L., Su, J. Z., Peng, X. Y., Shang, G. H., Han, D., Jin, J. Y., Liu, H. K., Du, Z. Y. et al. (2018). Different physiological roles of insulin receptors in mediating nutrient metabolism in zebrafish. American Journal of Physiology-Endocrinology and Metabolism 315, E38–E51.

Youson, J. H., Plisetskaya, E. M. and Leatherland, J. F. (1994). Concentrations of insulin and thyroid hormones in the serum of landlocked Sea lampreys (*Petromyzon marinus*) of three larval year classes, in larvae exposed to two temperature regimes, and in individuals during and after metamorphosis. General and Comparative Endocrinology 94, 294–304.

Zadik, Z., Chalew, S. A., Mccarter, R. J., Meistas, M. and Kowarski, A. A. (1985). The Influence of Age on the 24-Hour Integrated Concentration of Growth-Hormone in Normal Individuals. Journal of Clinical Endocrinology & Metabolism 60, 513–516.

Zoeller, R. T., Tan, S. W. and Tyl, R. W. (2007). General background on the hypothalamic-pituitary-thyroid (HPT) axis. Critical Reviews in Toxicology 37,11–53.

Zonneveld, C. and Kooijman, S. (1989). Application of a dynamic energy budget model to *Lymnaea stagnalis* (L). Functional Ecology 3, 269–278.

